# FSP1-mediated lipid droplet quality control prevents neutral lipid peroxidation and ferroptosis

**DOI:** 10.1101/2025.01.06.631537

**Authors:** Mike Lange, Michele Wölk, Cody E. Doubravsky, Joseph M. Hendricks, Shunji Kato, Yurika Otoki, Benjamin Styler, Kiyotaka Nakagawa, Maria Fedorova, James A. Olzmann

## Abstract

Lipid droplets (LDs) are organelles that store and supply lipids based on cellular needs. While mechanisms preventing oxidative damage to membrane phospholipids are established, the vulnerability of LD neutral lipids to peroxidation and protective mechanisms are unknown. Here, we identify LD-localized Ferroptosis Suppressor Protein 1 (FSP1) as a critical regulator that prevents neutral lipid peroxidation by recycling coenzyme Q10 (CoQ10) to its lipophilic antioxidant form. Lipidomics reveal that FSP1 loss leads to the accumulation of oxidized triacylglycerols and cholesteryl esters, and biochemical reconstitution of FSP1 with CoQ10 and NADH suppresses triacylglycerol peroxidation *in vitro*. Notably, polyunsaturated fatty acid (PUFA)-rich triacylglycerols enhance cancer cell sensitivity to FSP1 loss and inducing PUFA-rich LDs triggers triacylglycerol peroxidation and LD-initiated ferroptosis when FSP1 activity is impaired. These findings uncover the first LD lipid quality control pathway, wherein LD-localized FSP1 maintains neutral lipid integrity to prevent the buildup of oxidized lipids and induction of ferroptosis.

## INTRODUCTION

Lipid droplets (LDs) are the primary cellular organelle for lipid storage^1,2^. LD biogenesis is characterized by the biosynthesis of neutral lipids, such as triacylglycerols (TGs) and cholesteryl esters (CEs), that are deposited between the leaflets of the ER bilayer^1,3^ and ultimately phase separate and bud out into the cytoplasm. Structurally, LDs are unique when compared to other organelles. They lack a phospholipid bilayer and consist of a core of neutral lipids encased in a single phospholipid monolayer. While the core is devoid of proteins, regulatory proteins localize to the LD surface, influencing LD formation, organelle interactions, and stability^1,4^. Functionally, LDs serve as dynamic lipid storage depots, sequestering excess lipids and releasing lipids when needed for cellular processes^1^. Lipids stored in LDs can be mobilized for membrane biosynthesis, signaling molecule production, or energy generation through β-oxidation^1^. Moreover, LDs play a protective role by isolating lipids that could otherwise generate harmful species^1^, thereby safeguarding mitochondrial function^5^ and reducing ER stress^6^. Dysregulation of LD function can also impact the sensitivity of cells to the induction of ferroptosis^7–11^, an iron-dependent form of regulated cell death driven by oxidative damage to membrane phospholipids (i.e., lipid peroxidation)^12,13^.

Cells employ sophisticated quality control mechanisms and homeostatic pathways to preserve the integrity of biomolecules, including well-known pathways such as ubiquitin-dependent protein degradation^14^ and DNA damage repair^15^. Similarly, lipids are vulnerable to damage, such as oxidation, necessitating robust lipid quality control systems to prevent and counteract such damage^16^. In essence, ferroptosis represents a failure of lipid quality control, characterized by unchecked phospholipid peroxidation and consequent plasma membrane rupture^16^. To combat oxidative lipid damage and suppress ferroptosis, cells rely on specialized lipid quality control mechanisms. The glutathione-dependent peroxidase GPX4 plays a pivotal role by converting lipid peroxides into benign lipid alcohols^13,17^. In parallel, ferroptosis suppressor protein 1 (FSP1) functions by recycling lipophilic quinone radical trapping antioxidants, such as coenzyme Q10 (CoQ10)^18,19^ and vitamin K^20,21^, into their antioxidant forms. These antioxidants neutralize lipid peroxyl radicals, effectively halting the propagation of lipid peroxidation^16^.

Beyond the regulation of antioxidants, cells also modulate ferroptosis sensitivity by regulating the membrane lipid composition. Polyunsaturated fatty acids (PUFAs) are particularly prone to oxidation compared to monounsaturated fatty acids (MUFAs), and increasing the phospholipid MUFA:PUFA ratio reduces ferroptosis sensitivity across various contexts^22,23^. This shift can be achieved by supplementing cells with excess MUFAs (e.g., oleate)^24^ or by regulation of lipid metabolic enzymes, such as acyl-CoA synthetase (ACSL)-dependent fatty acid activation^25–27^ or membrane bound O-acyltransferase (MBOAT) family-dependent fatty acid incorporation into phospholipids)^28–30^. Furthermore, LDs have also been implicated in the regulation of the phospholipid MUFA:PUFA composition. By sequestering PUFAs in stored TG, LDs restrict their incorporation into membrane phospholipids, thereby decreasing membrane oxidizability and reducing cellular susceptibility to ferroptosis^7–9,11^.

While the mechanisms underlying membrane phospholipid peroxidation have been extensively studied, much less is known about the susceptibility of LD residing neutral lipids to peroxidation. Whether neutral lipids within LDs are prone to oxidative damage and what cellular mechanisms exist to protect them remain largely unexplored. In this study, we investigate the role of FSP1 in protecting stored neutral lipids from oxidative damage using a combination of lipidomic analyses in genetically engineered cells and biochemically reconstituted systems. Our results reveal that FSP1, together with its substrate CoQ10, acts at LDs to safeguard stored neutral lipids. This protection prevents the accumulation of TG and CE lipid peroxidation products, thereby averting an LD-initiated ferroptosis pathway. These findings uncover the first LD quality control pathway dedicated to protecting neutral lipids from oxidation and highlight a context-specific role for FSP1 in preventing LD-initiated ferroptosis.

## RESULTS

### FSP1 is required to maintain PUFA-containing di- and triacylglycerols

FSP1 suppresses lipid peroxidation and ferroptosis by recycling lipophilic antioxidants^18,19^. However, the specific lipids protected by FSP1 remain poorly characterized. To advance our understanding of the cellular role of FSP1, we performed untargeted lipidomics to analyze the lipid profiles of FSP1 knockout (KO) cells and KO cells expressing doxycycline (DOX)-inducible wild-type (WT) or catalytically inactive E156A mutant FSP1 (**Fig. 1**, **Extended Data Fig. S1A**). FSP1 has been established as a key ferroptosis resistance factor in U-2 OS osteosarcoma cells^19^. However, the basal levels of neutral lipids and LDs in this cell type are low, limiting the ability to assess the potential role of FSP1 in the regulation and oxidation of stored neutral lipids. To address this, we adapted our cell culture conditions and stimulated neutral lipid biosynthesis and LD biogenesis by supplementing the growth medium with a fatty acid mixture containing oleate (FA 18:1) and arachidonate (FA 20:4) (3:1 ratio) (**Extended Data Fig. S1B,C**).

**Figure 1.**
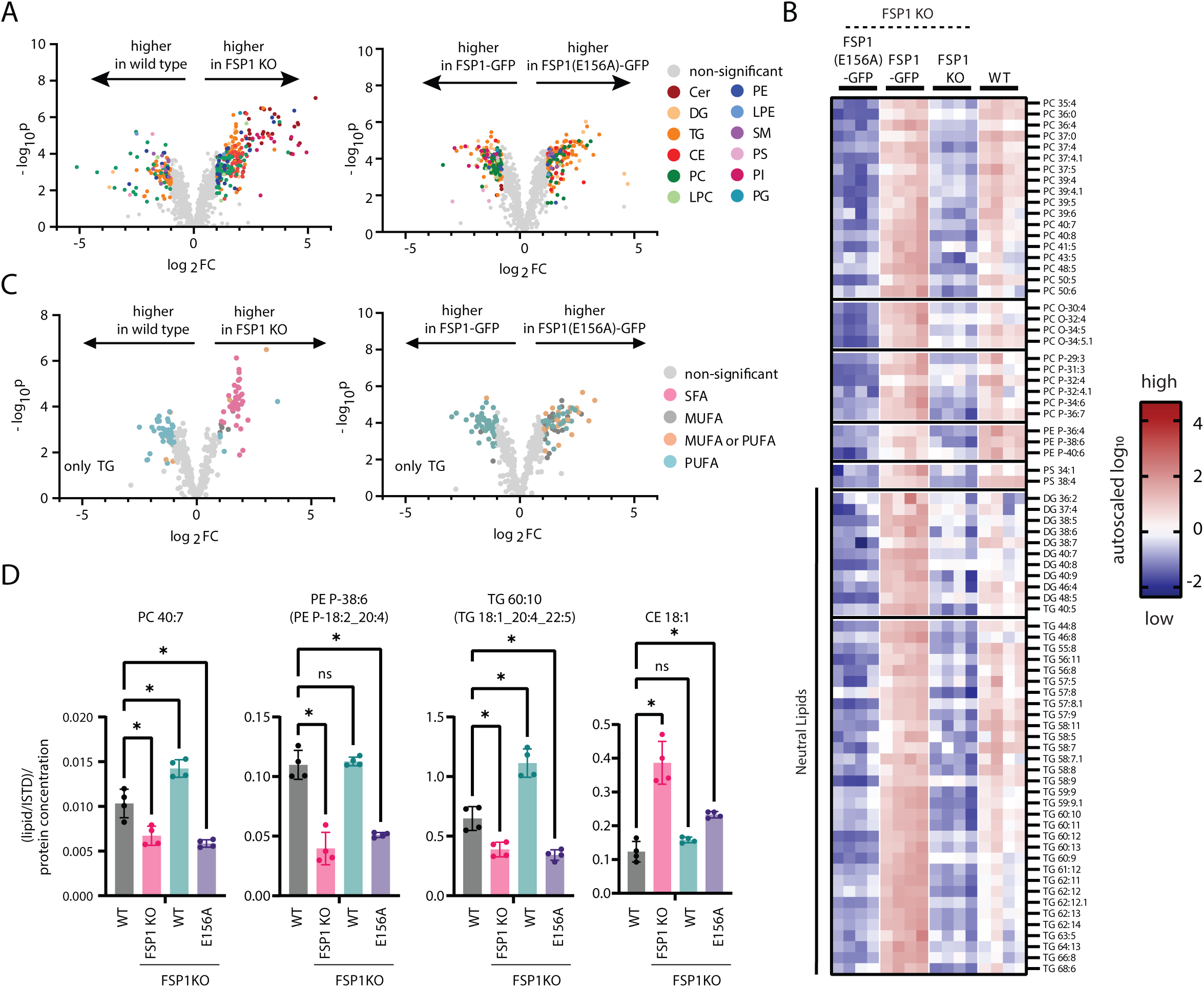
Loss of FSP1 results in lipid remodeling and a reduction in PUFA-containing glycerolipids. **A:** Volcano plots of untargeted lipidomics data comparing global lipidomes of wild type vs FSP1 KO and FSP1-GFP vs FSP1(E156A)-GFP re-expression constructs. Lipid classes are colored according to legend. Cer = ceramide, DG = diacylglcyerol, TG = triacylglycerol, CE = cholesteryl ester, PC = phosphatidylcholine, LPC = lysophosphatidylcholine, PE = phosphatidylethanolamine, LPE = lysophosphatidylethanolamine, SM = sphingomyelin, PS = phosphatidylserine, PI = phosphatidylinositol, PG = phosphatidylglycerol. (data was autoscaled and log10 transformed, thresholds for significant regulation are fold change > 2, p < 0.05 (FDR corrected)). **B:** Heatmap displaying a cluster of lipids that is protected by catalytically active FSP1 as identified by hierarchical clustering in Figure S1E. ((data was autoscaled and log10 transformed). **C:** Volcano plots of untargeted lipidomics data comparing triacylglycerol lipids of wild type vs FSP1 KO and FSP1-GFP vs FSP1(E156A)-GFP re-expression constructs. Triacylglycerol unsaturation is colored according to legend. SFA = saturated fatty acid containing (total double bonds = 0), MUFA = monounsaturated fatty acid containing (total double bonds = 1), PUFA = polyunsaturated fatty acid containing (total double bonds ≥ 4), MUFA or PUFA (total double bonds = 2 or 3). (data was autoscaled and log10 transformed, thresholds for significant regulation are fold change > 2, p < 0.05 (FDR corrected)). **D:** Selected lipids from FSP1 protected lipid cluster. (raw data is plotted, statistical significance is assessed by One-way ANOVA, * = p < 0.05).

Lipidomics revealed extensive alterations in lipid abundance across the experimental conditions (**Fig 1A**, **Extended Data Fig S1D**). Unbiased hierarchical clustering identified a subset of lipids that were preserved, or protected, by active FSP1 (**Fig 1B**, **Extended Data Fig S1D**). The levels of the lipids within this cluster were markedly reduced in FSP1 KO cells, and this phenotype was reversed by expression of WT FSP1 but not the catalytically inactive E156A mutant FSP1 (**Fig. 1B**, representative examples in **Fig 1D**). This cluster of lipids included highly unsaturated phospholipids, such as phosphatidylcholine (PC), phosphatidylethanolamine (PE), and phosphatidylserine (PS), in line with FSP1’s membrane protective function as well as many neutral lipids, including diacylglycerols (DGs) and TGs indicating an additional LD protective function (**Fig. 1B**). Indeed, FSP1 KO cells exhibited increased abundances of TGs containing saturated fatty acids and a concomitant reduction in PUFA-containing TG species (**Fig. 1A-C**) as well as other PUFA-containing phospholipids (**Extended Data Fig. S1E**). This indicates a key regulatory role of FSP1 in maintaining levels of unsaturated glycerolipids that are commonly found in LDs (**Fig. 1A-C**).

In addition to the lipids that decreased upon FSP1 KO, a subset of lipids showed increased levels. Notably, CEs were elevated in FSP1 KO cells, a change that was reversed by the expression of WT FSP1 and partially reversed by the catalytically inactive E156A mutant FSP1 (**Fig. 1A,D, Extended Data Fig. S1E**). CEs have previously been implicated in preventing TG lipid peroxidation *in vitro*^31^, suggesting that their increase may reflect an adaptive compensatory mechanism, where they might act as sacrificial antioxidants to mitigate lipid oxidative damage similar to the cholesterol intermediate 7-dehydrocholesterol^32–34^.

### FSP1 is required to prevent oxidative damage to cellular PUFA-containing neutral lipids

A striking finding from our lipidomics analyses is the critical role of FSP1 in the maintenance of PUFA-containing TG lipids (**Fig. 1**). One potential explanation for the observed reduction in PUFA-containing TGs in FSP1 KO cells is that they are being oxidized and thus no longer detected. Given the immense complexity of lipid species and their oxidized derivatives, we employed a semi-targeted liquid chromatography-tandem mass spectrometry (LC-MS/MS) approach for sample-specific epilipidome profiling^20,32,35,36^. This method involves identifying the most abundant PUFA-containing lipids in each sample and lipids specifically regulated by FSP1 inactivation, followed by predicting their potential oxidized species *in silico*, thereby generating refined inclusion lists for semi-targeted data-dependent acquisition during LC-MS/MS analysis^20,32,35,36^. Using this approach, we detected numerous oxidized neutral lipids that selectively accumulated in FSP1 KO cells, including oxidized TGs and CEs (**Fig. 2A,B**). These findings represent the first evidence that implicate FSP1 in protecting neutral lipids from oxidative damage.

**Figure 2.**
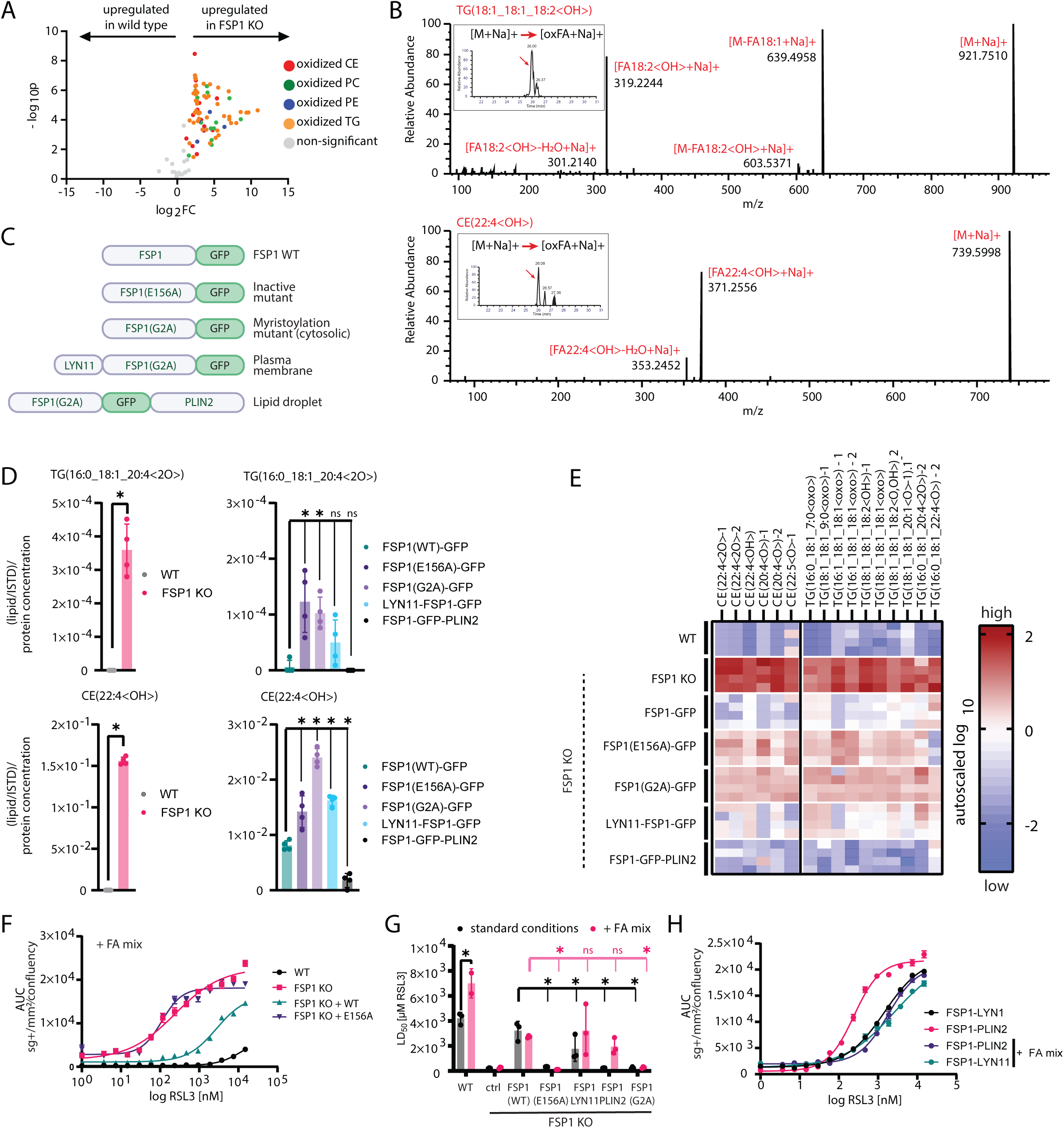
Lipid droplet-localized FSP1 prevents neutral lipid peroxidation and ferroptosis. **A:** Volcano plots of untargeted epi-lipidomics data comparing oxidized lipids of wild type vs FSP1 KO. Oxidized lipid class is colored according to legend. CE = cholesteryl ester, PC = phosphatidylcholine, PE = phosphatidylethanolamine, TG = triacylglycerol. (data was autoscaled and log10 transformed, thresholds for significant regulation are fold change > 2, p < 0.05 (FDR corrected)). **B:** Example MS/MS spectra from parallel reaction monitoring (PRM) based analysis of sodiated ions of oxidized TG and CE displaying informative fragments necessary for oxidized TG and CE identification. Inlet shows PRM based quantification of targeted transition (applied mass accuracy for extracted ion chromatogram generation = 5 ppm). **C:** Cartoon description of constructs re-expressed in FSP1 KO cells to study the role of FSP1 catalytic activity and subcellular localization. **D:** Selected oxidized lipids significantly elevated by FSP1 KO (raw data is plotted, statistical significance is assessed by One-way ANOVA, * = p < 0.05, ns = not significant). **E:** Heatmap displaying only oxidized triacylglycerol and oxidized cholesteryl ester lipids extracted from a cluster of oxidized lipids that is protected by catalytically active FSP1 as identified by hierarchical clustering in Figure S2E. (data was autoscaled and log10 transformed). **F:** Cell death sensitivity to RSL3 assessed by time resolved fluorescence microscopy (incucyte) with cell death indicator sytox green over the course of 24 hours. Number of sytox green positive objects per image equals the number of dead cells and is normalized to cell area per image expressed as confluency. sg+ = sytox green positive objects **G:** LD50 values for RSL3 in different cell lines under standard conditions or lipid droplet inducing conditions is determined by time resolved fluorescence microscopy (incucyte) using the cell death indicator sytox green. LD50 are calculated as the infliction point from the RSL3 dose-response curve under each condition. FA18:1 = oleate, FA20:4 = arachidonate. (statistical significance is assessed by One-way ANOVA, * = p < 0.05, ns = not significant). **H:** Cell death sensitivity to RSL3 assessed by time resolved fluorescence microscopy (incucyte) with cell death indicator sytox green over the course of 24 hours. Number of sytox green positive objects per image equals the number of dead cells and is normalized to cell area per image expressed as confluency. sg+ = sytox green positive objects.

FSP1 localizes to multiple organelles, including LDs and the plasma membrane^18,19^, raising the possibility that FSP1 influences oxidative damage to distinct sets of lipids at these discrete subcellular sites. FSP1’s dual localization raises the possibilities that the accumulation of oxidized neutral lipids induced by FSP1 inactivation may be due to 1) loss of local action of FSP1 at LDs actively protecting neutral lipids or 2) loss of distal action of FSP1 preventing oxidative damage to plasma membrane phospholipids whose damaged PUFAs are subsequently incorporated into LDs for detoxification. To investigate which possibility is prevalent, we analyzed cell lines expressing differentially targeted FSP1 constructs^19^ (**Fig. 2C, Extended Data Fig S2A**). Alongside WT and catalytically inactive E156A mutant FSP1, we included a myristoylation-defective mutant (G2A) which impairs membrane and LD recruitment, as well as constructs targeting FSP1 selectively to the plasma membrane (LYN11 fusion) or LDs (PLIN2 fusion). This experimental system allows us to dissect FSP1’s roles across subcellular compartments. Expression of WT FSP1 significantly suppressed the loss of PUFA-containing glycerolipids and the oxidation of TGs and CEs in FSP1 KO cells (**Fig. 2D,E, Extended Data Fig S2B,C**). In contrast, the E156A and G2A mutants exhibited diminished protective effects, highlighting the importance of FSP1’s enzymatic activity and membrane or LD association (**Fig. 2D,E**). Remarkably, FSP1 targeted specifically to LDs, but not to the plasma membrane, strongly suppressed TG and CE oxidation (**Fig. 2D,E**). These results indicate that LD-localized FSP1 is sufficient to protect neutral lipids from oxidative damage, suggesting that FSP1 acts directly at LDs to preserve the integrity of stored TGs and CEs.

FSP1 is well established as a ferroptosis suppressor^18,19^, but its specific roles in LD function and the cellular consequences of oxidized neutral lipids are unknown. WT FSP1, but not the catalytically inactive E156A mutant, conferred resistance to RSL3-induced ferroptosis in FSP1 KO cells grown under both standard conditions (**Extended Data Fig. 3A,B**) or under fatty acid supplemented conditions to induce LD biogenesis (**Fig. 2F**). Our previously published results indicated that plasma membrane targeted FSP1 was sufficient to suppress ferroptosis, but that LD-targeted FSP1 had no effect of ferroptosis resistance^19^. However, those assays were performed under standard growth conditions^19^ in which U-2 OS cells have low TG levels and few LDs. To investigate the impact of LD-localized FSP1 and neutral lipid peroxidation on ferroptosis, we performed dose-response analyses using increasing concentrations of RSL3 in our panel of FSP1-targeted cell lines pretreated with fatty acids to induce LDs. As anticipated, WT FSP1 and plasma membrane-targeted FSP1 effectively prevented RSL3-induced cell death under both standard and fatty acid-supplemented conditions, while the E156A and G2A mutants failed to confer protection (**Fig. 2G, Extended Data Fig S3B**). Interestingly, although LD-targeted FSP1 exhibited no protective effect under standard growth conditions, consistent with our previous results^19^, it strongly suppressed ferroptosis under fatty acid-supplemented conditions in which cells contain LDs (**Fig. 2G,H**). These findings highlight the critical and context-specific role of LD-localized FSP1 in protecting stored neutral lipids from oxidative damage, thereby suppressing ferroptosis under conditions characterized by elevated LD levels.

### CoQ10 is present in lipid droplets isolated from cells

FSP1 functions as an NAD(P)H oxidoreductase for quinone antioxidants, including CoQ10^18,19^ and vitamin K^20,21^, generating their reduced antioxidant forms which act as radical trapping antioxidants to inhibit the propagation of lipid peroxidation^18–20^. To investigate the presence of these molecules in intracellular LDs, we established a targeted LC-MS/MS method to detect derivatives of coenzyme Q, vitamin K, and vitamin E in LD-enriched buoyant fractions isolated from wild type U-2 OS cells under fatty acid stimulated conditions. These fractions were highly enriched with TG and CE (**Fig. 3A**). Among the quinone antioxidants, only CoQ10 (and small levels of CoQ9) was detected in the LD fractions (**Fig. 3B**). To gain quantitative insights, we developed an absolute quantification method by spiking isotopically labeled CoQ10, enabling precise measurement of oxidized CoQ10, reduced CoQ10H2, and total CoQ10 levels in LD-enriched fractions. This analysis confirmed that FSP1’s known substrate, CoQ10, is present in LDs, with approximately 60% in its antioxidant form, CoQ10H2. This approach also indicated that CoQ10 is present at approximately 1 molecule per 2,000 molecules of neutral lipids (**Fig 3C**). Together, these findings suggest that FSP1 acts via local recycling of CoQ10 to prevent neutral lipid peroxidation and maintain the antioxidant capacity in these organelles.

**Figure 3.**
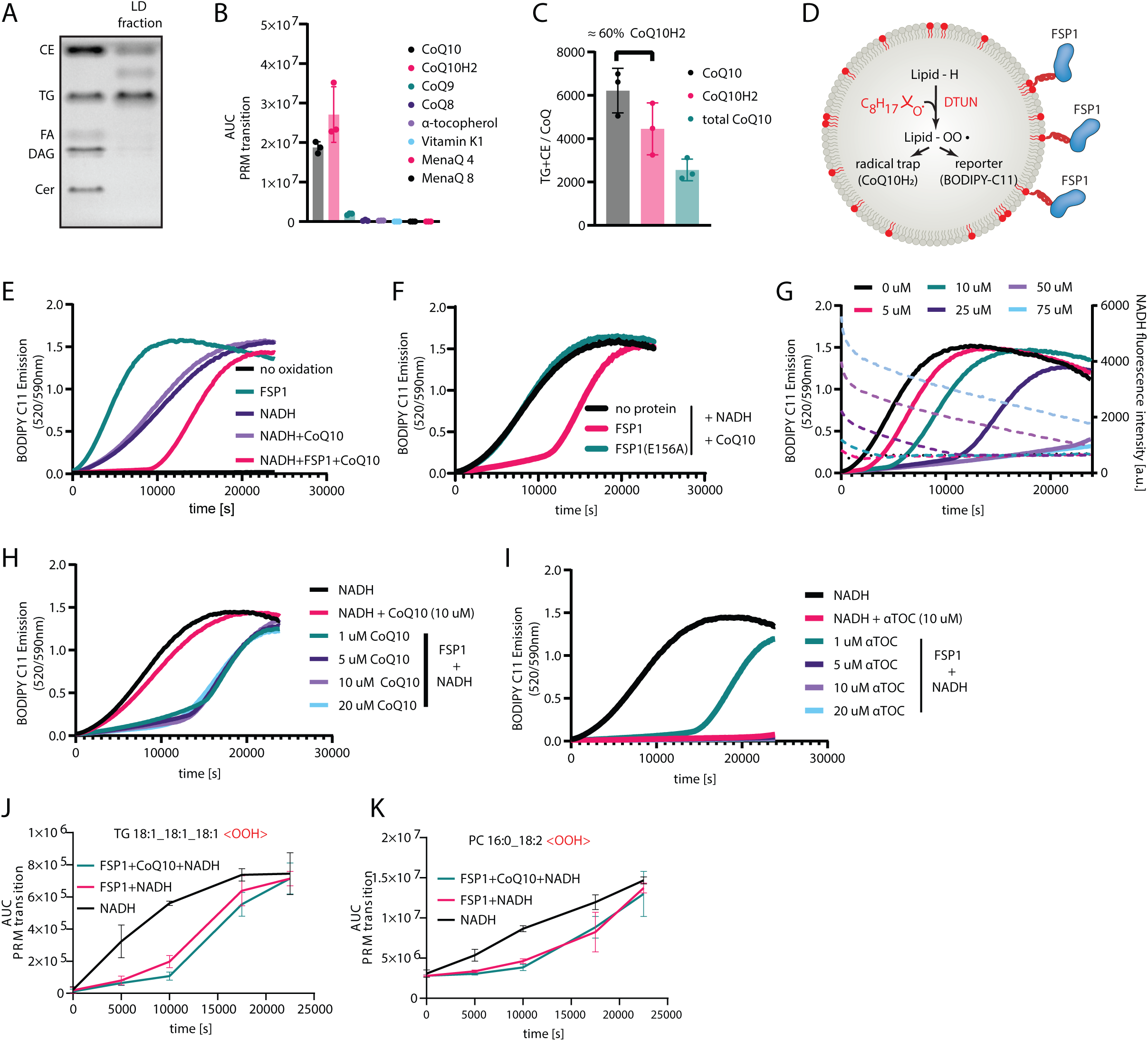
FSP1 reduction of CoQ10 suppresses triacylglycerol peroxidation in artificial lipid droplets. **A:** Thin layer chromatogram of lipids extracted from lipid droplet fractions. **B:** Parallel reaction monitoring used for relative quantification of known FSP1 substrates in lipid droplet fractions generated by density gradient centrifugation from cell lysates. **C:** Absolute quantification of CoQ10 levels by parallel reaction monitoring in lipid droplet fractions spiked with an isotopically labeled CoQ10 internal standard. Levels of CoQ10 were normalized to total amounts of TG and CE lipids quantified via thin layer chromatography. **D:** Cartoon describing the fluorescence enabled autooxidation assay (FENIX) adapted into artificial lipid droplets after recruitment of recombinant FSP1. **E:** FENIX assay in artificial lipid droplets with various substrate and enzyme compositions. **F:** FENIX assay in artificial lipid droplets in the presence and absence of catalytically active FSP1 or the catalytically inactive FSP1(E156A) mutant. **G:** FENIX assay in artificial lipid droplets in the presence of 100 nM FSP1, 10 uM CoQ10, 1 mM total lipid and and varying concentrations of NADH. NADH consumption throughout the FENIX assay was assessed via NADH autofluorescence within the same experiment. **H/I:** FENIX assay in artificial lipid droplets in the presence of increasing concentrations of **(H)** CoQ10 or **(I)** α-tocopherol. **J/K:** Parallel reaction monitoring based quantification of TG and PC oxidation products during peroxidation of artificial lipid droplets.

### FSP1 suppresses oxidative damage to TG in a biochemically reconstituted system

To examine the mechanisms of FSP1 in suppressing lipid peroxidation, we used biochemically reconstituted approaches. Recombinant His-tagged WT and E156A mutant FSP1 were purified from bacteria (**Extended Data Fig. S4A**). Purified WT FSP1, but not the E156A mutant, displays the expected activity and consumes NADH to in the presence of the soluble CoQ10 analogue CoQ1 and reduces the soluble and fluorescent CoQ10 analogue CoQ1-coumarin to its quinol form in the presence of NADH (**Extended Data Fig. S4B,C**). To determine if recombinant FSP1 is able to suppress lipid peroxidation using CoQ10, we generated liposomes containing trace amounts of nickel conjugated phospholipids to recruit His-tagged FSP1 to the liposome surface (**Extended Data Fig. S4D**). Employing the FENIX (fluorescence-enabled inhibited autoxidation) assay^37^, lipid peroxidation was triggered using the lipid radical initiator di-tert-undecyl hyponitrite (DTUN) and lipid peroxidation propagation was followed by BODIPY-C11 fluorescence (**Extended Data Fig. S4E**). In this reconstituted liposome system, WT FSP1, but not the E156A mutant, prevented lipid peroxidation when its substrates NADH and CoQ10 were present (**Extended Data Fig. S4E,F**). FSP1 also efficiently suppressed lipid peroxidation in the presence of NADH and the vitamin E derivative α-tocopherol (**Extended Data Fig. S4G**). These findings support the mechanism of FSP1 in suppressing membrane phospholipid peroxidation.

To test the ability of FSP1 to directly protect TG from oxidative damage in LDs, we developed a modified FENIX assay that uses artificial LDs generated *in vitro* (**Fig. 3D, Extended Data Fig. S5A**). These artificial LDs mimic cellular LDs, exhibiting a uniform size of ∼290 nm in diameter (**Extended Data Fig. S5B,C**) and containing a core of TG (triolein) surrounded by a phospholipid monolayer as indicated by thin layer chromatography analysis (**Extended Data Fig. S5D,E**) and the fluorescence emission spectrum of the solvatochromatic dye nile red (**Extended Data Fig. S5F**). Similar to our liposome assays, His-tagged FSP1 was recruited to the LD surface using nickel conjugated phospholipids (**Extended Data Fig. S5H**). Following the initiation of lipid peroxidation with DTUN, we observed unrestricted lipid peroxidation based on the oxidation of BODIPY-C11(**Fig. 3E**). The addition of NADH marginally reduced the rate of lipid peroxidation (**Fig. 3E**), potentially by quenching some of the radical species. However, NADH did not fully prevent lipid peroxidation indicated by the absence of an initial lag phase. In contrast the addition of FSP1, together with CoQ10 and NADH, resulted in full inhibition of lipid peroxidation propagation evident by an extended lag phase (**Fig. 3E, Extended Data Fig. S5I**). As expected, the E156A mutant FSP1 had no effect on the kinetics of lipid peroxidation (**Fig. 3F, Extended Data Fig. S5J**). In the presence of FSP1, NADH, and CoQ10, the lag phase continued for approximately 10,000 seconds and then unrestricted lipid peroxidation was observed (**Fig. 3E,F**). These findings suggested that a component within the reaction was being consumed. Indeed, the addition of increasing concentrations of NADH resulted in an extension in the lag phase followed by lipid peroxidation upon complete consumption of NADH (**Fig. 3G**). In contrast, addition of higher amounts of CoQ10 did not affect the reaction (**Fig. 3H**), indicating that 1 µM CoQ10 is sufficient to drive maximal suppression of lipid peroxidation in this assay. FSP1 was also able to suppress lipid peroxidation in the presence of NADH together with the vitamin E derivative α-tocopherol or the vitamin K derivative menaquinone-4 (**Fig. 3I, Extended Data Fig 5K**). Given that we were unable to detect *α*-tocopherol or menaquinone-4 in LDs purified from our cells, it is likely that FSP1 uses CoQ10 to prevent neutral lipid peroxidation in LDs. However, there may be cell types or metabolic conditions in which α-tocopherol or menaquinone-4 are present in LDs, and FSP1 could employ these other quinone antioxidants to maintain LD lipid quality.

*In vitro* generated artificial LDs contain both phospholipids and TG. An indirect lipid peroxidation indicator such as BODIPY C11 is unable to discern which lipids are being oxidized. Indeed, FSP1 may affect the oxidation of stored TG, the phospholipids in the bounding monolayer, or both. To determine which lipids are oxidized and the impact of FSP1, we performed semi-targeted oxidized epilipidomics analyses. As expected, due to the simple lipid composition and peroxidation conditions, we detected only a small number of oxidized lipids. Our findings revealed that FSP1, together with CoQ10 and NADH, suppresses the oxidation of both TG (**Fig. 3J, Extended Data Fig. S6A-E**) and phospholipids (**Fig. 3K, Extended Data Fig. S6F-I**). Thus, our findings in cellular and reconstituted systems indicate that FSP1 acts as a CoQ10 oxidoreductase to prevent the oxidative damage to neutral lipids stored in LDs.

### PUFA supplementation generates PUFA-rich LDs that are prone to peroxidation

To understand the physiological importance of FSP1’s LD protective function, we examined correlations of FSP1 essentiality and cellular lipid levels across a broad panel of cancer cell lines. Analysis of FSP1 dependency in DepMap revealed a correlation between FSP1 essentiality and the amount of PUFA-containing TG across 649 cancer cell lines (**Fig 4A,B**). Although, FSP1 is not essential in most cell types, FSP1 is required for the viability of a subset of cell types with high PUFA TG (**Fig 4A,B**). PUFA-containing TG levels (e.g., TG 56:6) correlate with the essentiality for many genes related to ferroptosis, including genes involved in glutathione synthesis, selenoprotein biosynthesis, fatty acid metabolism, and antioxidant defense (e.g., *GPX4* and *FSP1/AIFM2*) (**Fig 4C**). These data indicate a correlation between PUFA-containing TG and ferroptosis sensitivity, which may reflect the overall cellular lipid unsaturation environment or a vulnerability to ferroptosis conferred by the unsaturation of TG.

**Figure 4.**
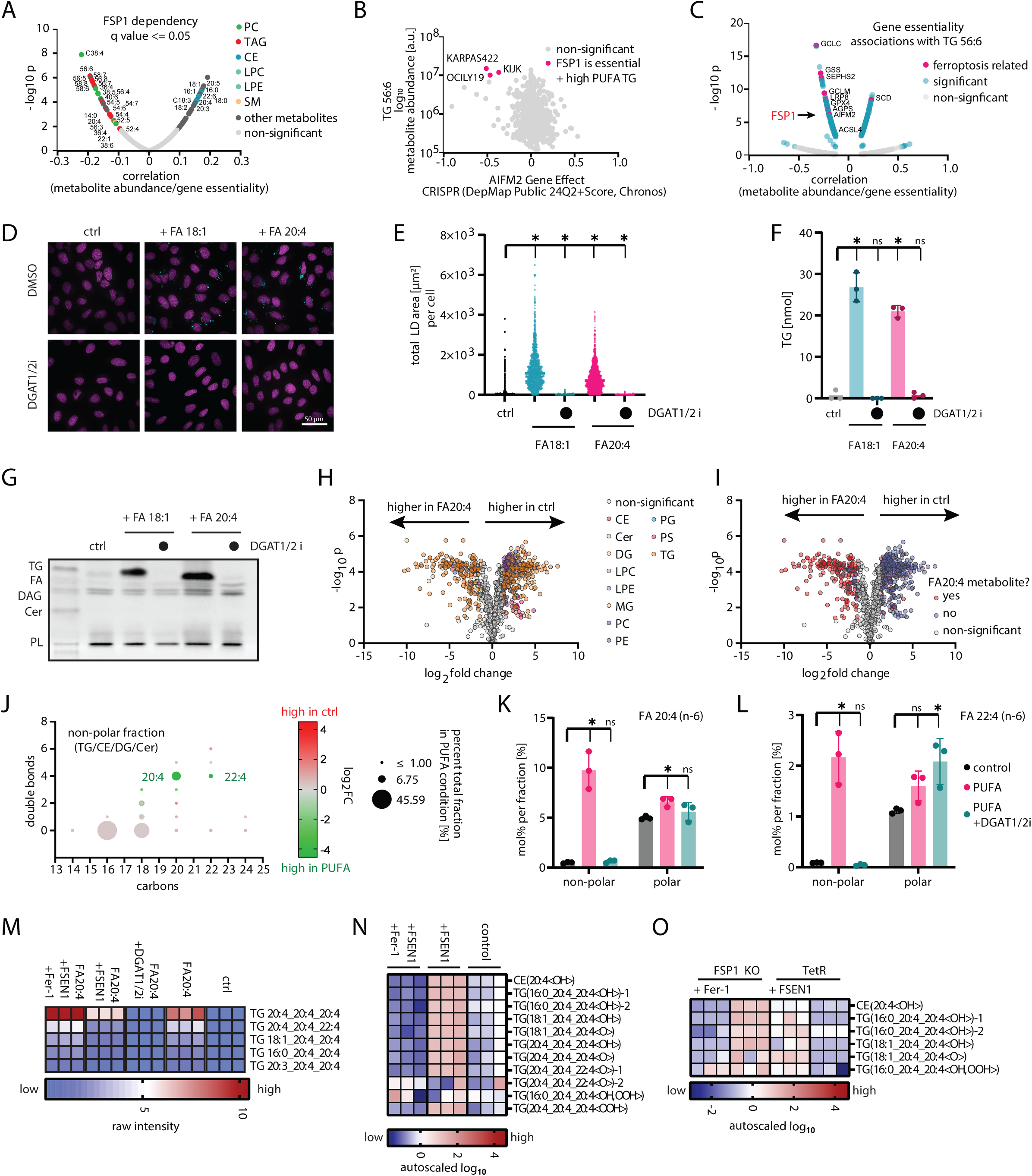
FSP1 prevents the peroxidation of PUFA-rich lipid droplets. **A:** Mining the cancer dependency portal DepMap for correlations of FSP1 essentiality with metabolite abundance indicates FSP1 essentiality correlates with elevated levels of PUFA-containing TG and PUFA-containing PC lipids. **B:** Example correlation of TG 56:6 with FSP1 essentiality across 927 cancer cell lines. Each dot represents one cell line. **C:** Association of relative abundance of TG 56:6 with gene essentiality indicates anti-ferroptotic genes correlate highly with PUFA-TG levels. **D:** Lipid droplet quantification in U-2 OS cells treated with 200 µM oleate (FA 18:1) or arachidonate (FA 20:4) for 24 hours in the presence and absence of DGAT1 (15 μM A-922500) and DGAT2 (10 μM PF-06424439) inhibitors. Lipid droplets were stained with BODIPY 493/503 and nuclei with Hoechst. **E:** Lipid droplets were imaged and lipid droplet area for each cell was quantified using Harmony data analysis software (Revvity). Each dot represents the lipid droplet area in an individual cell. (statistical significance is assessed by One-way ANOVA, * = p < 0.05, ns = not significant). **F:** Quantification of triacylglycerol levels via thin layer chromatography. (statistical significance is assessed by One-way ANOVA, * = p < 0.05, ns = not significant). **G:** Example of a thin layer chromatogram used for quantification in Fig 4F. **H:** LC-MS/MS based untargeted lipidomics comparing cells in the presence and absence of 200 µM FA 20:4 for 24 hours. Lipid classes are colored according to legend. Cer = ceramide, DG = diacylglcyerol, TG = triacylglycerol, CE = cholesteryl ester, PC = phosphatidylcholine, LPC = lysophosphatidylcholine, PE = phosphatidylethanolamine, LPE = lysophosphatidylethanolamine, SM = sphingomyelin, PS = phosphatidylserine, PI = phosphatidylinositol, PG = phosphatidylglycerol. (data was autoscaled and log10 transformed, thresholds for significant regulation are fold change > 2, p < 0.05 (FDR corrected)). **I:** Same data as displayed in Fig 4H. Significantly regulated lipids are colored based on the presence or absence of FA20:4 metabolites. A lipid was deemed a putative FA20:4 metabolite when either of its fatty acids had more than 20 carbons with more than 4 double bonds. **J:** Bubble plot showing absolute quantities and regulation of fatty acids in the non-polar lipid fraction extracted from U-2 OS cells treated with 200 µM FA20:4 for 24 hours. For quantification, lipid extracts were separated into non-polar lipid fraction using solid phase extraction and esterified fatty acids were transmethylated and quantified using gas chromatography-flame ionization detection. **K:** Mol% of arachidonate (FA20:4 (n-6)) in non-polar and polar lipid fractions. (statistical significance is assessed by One-way ANOVA, * = p < 0.05, ns = not significant). **L:** Mol% of adrenate (FA22:4 (n-6)) in non-polar and polar lipid fractions. (statistical significance is assessed by One-way ANOVA, * = p < 0.05, ns = not significant). **M:** Highest intense triacylglycerol lipids in cells treated with 200 µM FA20:4 for 24 hours quantified by untargeted lipidomics. **N:** Semi-targeted epi-lipidomics analysis of non-polar lipids in wild type U-2 OS cells treated with 200 µM FA20:4 for 24 hours. **O:** Semi-targeted epi-lipidomics analysis of non-polar lipids control and FSP1 KO U-2 OS cells treated with 20 µM FA20:4 for 24 hours.

To explore the possibility that PUFA-rich TG confers ferroptosis sensitivity and represents a vulnerability to the loss of FSP1, we developed a cell-based system in which we generate high amounts of PUFA-rich TG. We hypothesized that FSP1 inactivation in this background would preferentially lead to LD peroxidation. Incubation of U-2 OS cells with the MUFA oleate (FA 18:1) or the PUFA arachidonate (FA 20:4) induced comparable biosynthesis of TG and LDs, which were both dependent on the diacylglycerol acyltransferases (DGAT1/2) (**Fig. 4D-F**). Incubation with arachidonate altered the cellular lipid landscape, with the majority of changes being in the amounts of TG lipids (**Fig. 4G,H, Extended Data Fig. S7**). Analysis of TG lipid changes also revealed an overwhelming increase in TG lipids containing arachidonate or putative arachidonate elongation and desaturation products (**Fig. 4I**), which was dependent upon DGAT1/2 (**Extended Data Fig. S7**). A small number of phospholipids were also upregulated under arachidonate supplemented conditions, raising the possibility that arachidonate was also incorporated into the plasma membrane. The untargeted lipidomics approach utilized in this study does not provide quantitative information regarding fatty acid quantities within different lipid classes and only yields relative quantitative information for individual intact lipids. To directly quantify the fatty acid composition of LDs or membranes, we extracted total lipids from cells and isolated non-polar lipids (including TG, CE, DG) and polar lipids (including all phospholipid subclasses) (**Extended Data Fig. S8A,B**), and employed gas chromatography with flame ionization detection (GC-FID) to quantify the fatty acid composition in these fractions (**Fig. 4J-L, Extended Data Fig. S8C-F and S9**). This analysis indicated an increase in arachidonate and adrenic acid (22:4)-containing non-polar lipids (**Fig. 4J**) in the treated samples relative to control, with very little change observed in the polar lipids (i.e., phospholipids) (**Fig. S8F**). Importantly, DGAT inhibition blocked the increase in arachidonate and adrenic acid in non-polar lipids with little effect on the amount present in polar lipids (**Fig. 4K,L**). These data establish a cell-based system to generate intracellular LDs with high amounts of PUFA-containing TG.

We hypothesized that the PUFA-rich TG in arachidonate treated cells would be peroxidation prone and sensitive to the loss of FSP1 function. Indeed, untargeted lipidomics indicated that treatment with the FSP1 inhibitor FSEN1 reduced the highest abundant arachidonate-containing TG (**Fig. 4M, Fig. S8G**). Co-treatment with the radical trapping antioxidant ferrostatin (Fer-1) reversed this decrease (**Fig. 4M**), solidifying that the observed decrease of arachidonate-containing TG following FSP1 inhibition is peroxidation dependent. To test this possibility directly, we performed another semi-targeted epilipidomics analysis. Treatment with the FSP1 inhibitor FSEN1 resulted in the accumulation of oxidized arachidonate containing CE and TG lipids, which was blocked by co-treatment with Fer-1 indicating that oxidized TG and CE are generated by active lipid peroxidation (**Fig. 4N**). To gain more specificity on the role of FSP1 in preventing TG and CE oxidation and to exclude off-target effects of FSP1 inhibitors, we also performed a similar assay in FSP1 KO cells. FSP1 KO cells display heightened cell death sensitivity to arachidonate treatment, thus a lower arachidonate concentration had to be used to induce a PUFA TG phenotype, in order to maintain viability of FSP1 KO cells. Treatment of FSP1 KO cells with 20 μM arachidonate induced a similar upregulation of TGs containing arachidonate and putative arachidonate metabolites that could be blocked with DGAT1/2 inhibitors (**Fig S7C-H**). Within this system FSP1 KO similarly resulted in an accumulation of oxidized CE and TG lipids, which were suppressed by Fer-1 co-treatment (**Fig. 4O**). These data indicates that PUFA-containing neutral lipids are highly reliant upon FSP1 to prevent their peroxidation.

### FSP1-mediated neutral lipid quality control prevents LD-initiated ferroptosis

Our cellular system, utilizing arachidonate treatment to induce PUFA-rich TG-containing LDs, provides a controlled model to investigate the cellular consequences of neutral lipid peroxidation. Arachidonate treatment was well tolerated by control cells, with minimal cell death observed at concentrations up to 200 µM (**Fig. 5A**). In contrast, FSP1 KO cells exhibited heightened sensitivity to arachidonate, with cell death occurring at concentrations as low as 25 µM and substantial cell death observed at 200 µM (**Fig. 5A**). This arachidonate-induced cell death in FSP1 KO cells was reversed by co-treatment with the ferroptosis inhibitor and radical trapping antioxidant Fer-1 (**Fig. 5A**), underscoring the role of lipid peroxidation in this process and consistent with this cell death via ferroptosis. Similarly, control cells co-treated with the FSP1 inhibitor FSEN1 displayed increased sensitivity to arachidonate-induced cell death (**Fig. 5B, Extended Data Fig. S10A,B**). Notably, although treatment with the monounsaturated fatty acid oleate increased LDs similar to arachidonate treatment (**Fig. 4D-F**), it did not induce appreciable cell death and there were no significant differences between control and FSP1 KO cells (**Fig. 5C**). These findings identify a cell death that is induced by the peroxidation of PUFA-rich LDs even in the absence of targeting the GSH-GPX4 pathway, highlighting the critical role of FSP1 in protecting cells from ferroptosis via maintaining neutral lipid quality control.

**Figure 5.**
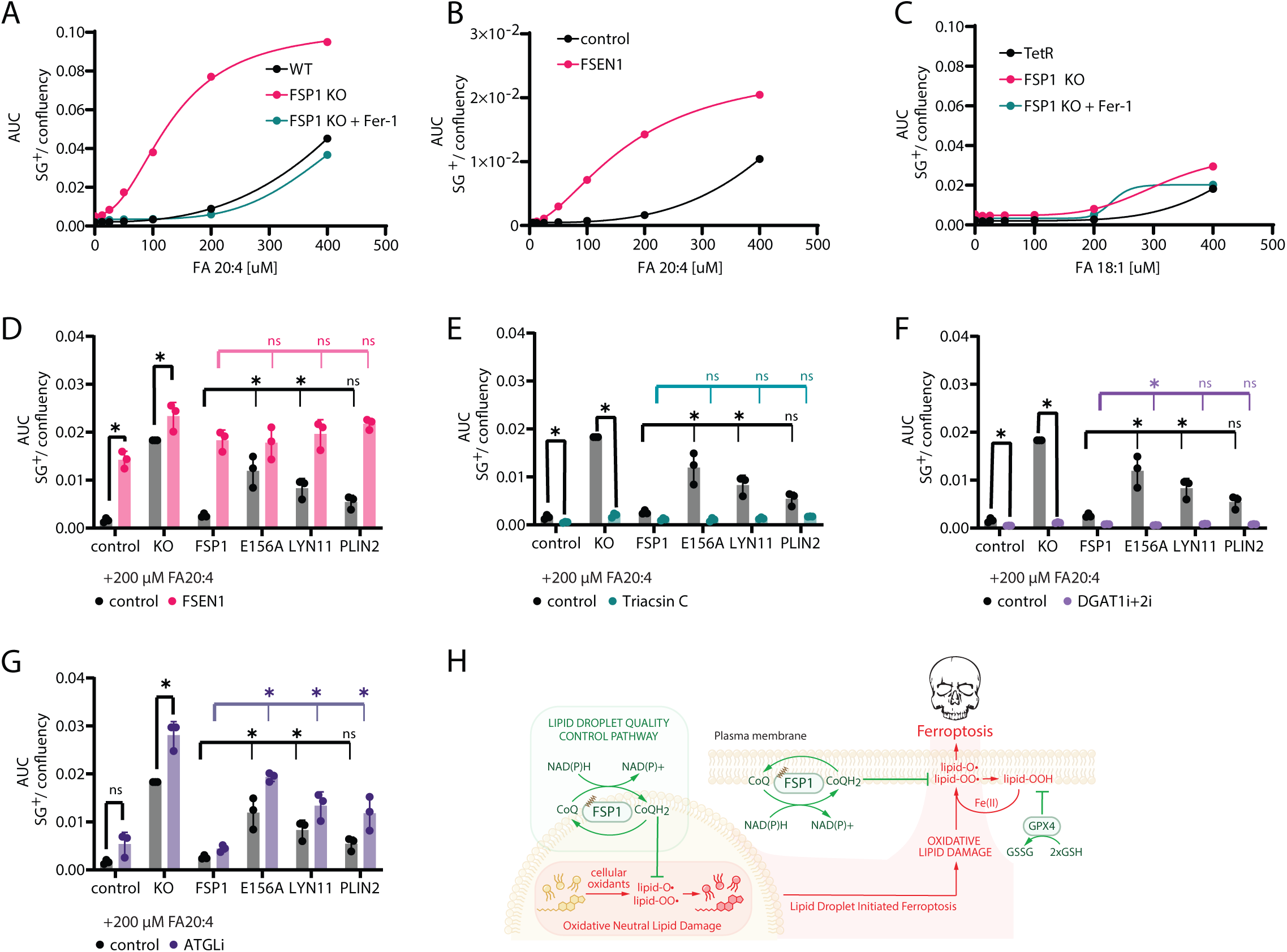
FSP1 suppresses lipid droplet-initiated ferroptosis. **A-C** : Cell death sensitivity to arachidonate (FA20:4) (panels **A,B**) or oleate (FA18:1) (panel **C**) treatment assessed by time resolved fluorescence microscopy (incucyte) with cell death indicator sytox green over the course of 24 hours. Number of sytox green positive objects per image equals the number of dead cells and is normalized to cell area per image expressed as confluency. sg+ = sytox green positive objects. **D-G:** Cell death quantified at 200 µM FA20:4 after 24 hours in the presence of FSP1 inhibitor (FSEN1 = cotreatment with 5 μM FSEN1), DGAT1/2 inhibitors (= cotreatment with 15 μM A-922500 and 10 μM PF-06424439), acyl-CoA synthetase inhibitor (Triacsin C = cotreatment with 1 μg\ml Triacsin C) or TG lipolysis inhibitor (ATGLi = cotreatment with 10 μM NG-497). (statistical significance is assessed by One-way ANOVA, * = p < 0.05, ns = not significant). **H:** Schematic depicting the proposed mechanism of LD protection by FSP1.

To investigate the significance of FSP1 activity and localization, we utilized our panel of cell lines expressing various FSP1 constructs. Expression of WT FSP1 strongly suppressed arachidonate-induced cell death in FSP1 KO cells, whereas this protective effect was abolished by the catalytically inactive E156A mutation (**Fig. 5D**). Both plasma membrane-targeted and LD-targeted FSP1 constructs partially mitigated arachidonate-induced cell death in FSP1 KO cells, although neither was as effective as WT FSP1, which localizes to multiple cellular compartments (**Fig. 5D**). The suppression of cell death by FSP1 was reversed by co-treatment with the FSP1 inhibitor FSEN1 (**Fig. 5D**), emphasizing the importance of FSP1 catalytic activity in protecting cells under these conditions. To explore the importance of TG and LDs, we treated cells with the inhibitor of acyl-CoA synthetase triacsin C or with inhibitors of DGAT enzymes. Both triacsin C and DGAT1/2 inhibitors effectively prevented arachidonate-induced cell death in all cell types, including FSP1 KO cells (**Fig. 5E,F, Extended Data Fig. S10C**). These findings indicate that the cytotoxic effects of arachidonate in FSP1 KO cells are dependent on its activation and incorporation into TG. In contrast, inhibition of ATGL, the rate-limiting enzyme in TG lipolysis, failed to prevent arachidonate-induced ferroptosis and slightly potentiated cell death in the FSP1 KO cells (**Fig. 5G, Extended Data Fig. S10C**). Some small molecule inhibitors can act as radical trapping antioxidants, leading to confusion in the field. We therefore used the FENIX assay to test if DGAT1/2 or ATGL inhibitors are radical trapping antioxidants preventing ferroptosis via off-target antioxidant activity. Importantly, in contrast to the established radical trapping antioxidant Fer-1, the utilized DGAT1/2 inhibitors and the human ATGL inhibitor (NG-497) had no radical trapping antioxidant activity and support an on-target mechanism of action. We did note that the mouse ATGL inhibitor Atglistatin exhibited some radical trapping antioxidant activity. While we did not use this inhibitor in our studies, we wanted to share the data as a cautionary note for the field. Together, these results highlight the pivotal role of FSP1 in preventing ferroptosis induced by the accumulation of oxidatively damaged TGs in LDs.

## DISCUSSION

In this study, we discover a critical role for FSP1 and CoQ10 in safeguarding neutral lipids against peroxidation (**Fig. 5H**). Loss of FSP1 led to the accumulation of oxidized neutral lipids and triggered ferroptosis initiated from LDs (**Fig. 5H**). These findings identify the first LD-specific lipid quality control pathway that protects stored lipids from peroxidation and support the emerging concept that ferroptosis can originate from distinct subcellular compartments depending on cellular context. In addition, the buildup of oxidized neutral lipids and the induction of ferroptosis initiated from PUFA-rich LDs upon FSP1 inhibition underscores the critical role of FSP1 in safeguarding neutral lipids from peroxidation. Indeed, this occurs in the absence of GPX4 inhibitors, indicating that GPX4 cannot compensate for FSP1 loss under these conditions and identifying a context in which FSP1 acts as the primary ferroptosis suppressor.

Our data support a model in which LD-localized FSP1 reduces CoQ10 to generate local antioxidants that prevent neutral lipid peroxidation. This is evidenced by the reduction of PUFA-containing TGs and CEs, and the concomitant increase in their oxidized counterparts, upon FSP1 KO. The direct role of FSP1 is underscored by the strong suppression of neutral lipid oxidation when FSP1 is selectively targeted to LDs through fusion with the LD protein PLIN2 and in biochemical reconstitution assays where purified FSP1 and CoQ10 prevent TG peroxidation. CoQ10 frequently functions as a co-antioxidant with α-tocopherol, quenching tocopheroxyl radicals generated following the action of α-tocopherol as radical trapping antioxidant. However, LDs, which we find to lack detectable α-tocopherol or vitamin K in the conditions analyzed in this study, appear to represent a context in which CoQ10 acts directly as the major radical scavenger. It is plausible that other lipophilic radical trapping antioxidants contribute to preventing neutral lipid peroxidation in specific tissues or under distinct metabolic conditions. How CoQ10 traffics to LDs remains an open question. STARD7 is a lipid transfer protein that mediates CoQ10 trafficking to the plasma membrane^38^. One possibility is that STARD7 or a similar lipid transfer protein could deliver CoQ10 to LDs. Given the ER origin of LDs, it is also possible that CoQ10 traffics directly from the ER into LDs.

The accumulation of oxidized neutral lipids likely impacts the structural and functional properties of LDs. Molecular dynamics simulations suggest that PUFA peroxidation within TG molecules increases their polarity, leading them to migrate from the neutral lipid core to the periphery of the LD, with the oxygen atoms protruding from between the LD headgroups^39^. Simulations of oxidation of phospholipids have yielded similar models in which the oxidized fatty acids bend towards the aqueous phase resulting in increased membrane permeability^40–44^. Structural changes resulting from oxidized fatty acids protruding between phospholipid headgroups would be expected to disrupt LD phospholipid packing, which is known to play important roles in LD protein targeting and association^1,4^. LDs would not be expected to rupture in the traditional manner of other organelles, since their lumen is made of neutral lipids rather than an aqueous compartment, however, the increased exposure of the hydrophobic oil core to the aqueous cytosol would likely disrupt LD morphology and alter protein association with LDs. Additionally, the surface exposure of TG or CE peroxides could facilitate the propagation of lipid peroxidation within the LD by facilitating Fenton-type reactions with iron.

Our findings expand the paradigm that ferroptosis can be triggered from various subcellular compartments, including LDs, ER, and lysosomes. For example, ferroptosis induction has been proposed to involve lipid peroxidation initially in the ER and at later stages at the plasma membrane^24,45,46^. Alternatively, lipid peroxidation has been observed within lysosomes prior to lipid peroxide accumulation in the ER^47^, suggesting that, in certain contexts, ferroptosis may originate in the lysosome and subsequently propagate to other organelles. In our system, LD-initiated ferroptosis occurs under conditions of high PUFA-TG content and impaired FSP1 function. DGAT inhibitors, which block TG synthesis and LD biogenesis, effectively prevented ferroptosis in this context, underscoring the critical role of TGs and LDs in initiating ferroptosis under these conditions. Oxidation of LDs may contribute to ferroptosis under other conditions, such as ferroptosis induced by co-treatment with imidazole ketone erastin and the oxidized form of vitamin C^10^. Truncated forms and hydroxyl derivatives of TGs have also been observed in dendritic cells in cancer and hypoxic trophoblasts cells^39,48,49^. Though the importance of the oxidized TG in these contexts remains to be determined. The mechanisms by which lipid peroxidation propagates from intracellular organelles such as LDs to the plasma membrane remain unclear. We ruled out lipolysis as a requirement, as ATGL inhibitors had no effect or slightly enhanced LD-initiated ferroptosis. It is possible that exposed TG peroxides at the LD surface could propagate oxidative damage by promoting iron oxidation or LD phospholipid peroxides could be transferred to other membranes at membrane contact sites. The ER’s connection with LDs is notable, as LDs can grow through Ostwald ripening, a process in which the ER acts as a conduit for TG trafficking that enables neutral lipid exchange between LDs^50^. This suggests a potential pathway for the transmission of lipid peroxidation from LD neutral lipids to ER phospholipids, which could then spread through vesicular trafficking or phospholipid transfer at membrane contact sites.

Our findings also imply a broader context for FSP1’s role in lipid quality control, with implications for physiology and disease pathogenesis. LDs play essential roles in metabolically active tissues, such as adipose tissue and the liver, where they regulate both local and systemic lipid metabolism. The duration of lipid storage within these tissues varies greatly, ranging from just a few hours to several years. During this time, maintaining lipid quality—particularly under stress—is critical to ensuring a dependable supply of undamaged lipids when the body needs energy. Notably, FSP1 is strongly expressed in brown adipose tissue^51^, a metabolically active tissue with high amounts of LDs and mitochondria. This raises intriguing questions about whether FSP1 also plays a critical role in maintaining lipid homeostasis and protecting against lipid peroxidation in energy-dense tissues. In addition, LDs are known to accumulate under various pathological conditions. For instance, neurons efflux fatty acids for sequestration within astrocytic LDs as a protective mechanism^52–57^, and recent findings suggest that these fatty acids may include fatty acid peroxides^55,56^. It remains an open question whether FSP1 plays a role in LD quality control in astrocytes by mitigating the spread of LD peroxidation. Similarly, ferroptosis has been implicated in the pathology of liver diseases characterized by LD accumulation and the buildup of PUFA-containing TGs, such as metabolic dysfunction-associated fatty liver disease (MAFLD) and metabolic-associated steatohepatitis (MASH)^58–60^. These conditions, which are closely tied to systemic metabolic dysregulation, highlight the potential importance of LD quality control mechanisms in protecting against ferroptosis-induced cellular damage.

In conclusion, our study highlights the susceptibility of neutral lipids to oxidation and identifies LD-localized FSP1 as a critical mediator of LD lipid quality control and the cellular defense against LD-initiated ferroptosis. Through its reduction of CoQ10, FSP1 maintains neutral lipid integrity, preventing cellular dysfunction and oxidative damage propagation. Our findings underscore the need for further research into neutral lipid peroxidation, FSP1-mediated LD quality control, and their roles in physiology and the pathogenesis of LD-associated and oxidative stress-driven diseases.

## AUTHOR INFORMATION

Correspondence and requests for materials should be addressed to J.A.O. and M.L.

## ACKNOWLEDGEMENTS

This research was supported by a grant from the National Institutes of Health to J.A.O. (R01GM112948) and Bakar Fellows Spark Award to J.A.O. Work in the Fedorova lab is supported by ‘‘Sonderzuweisung zur Unterstützung profilbestimmender Struktureinheiten’’ by the SMWK to TUD, TG70 by Sächsische Aufbaubank and SMWK, the measure is co-financed with tax funds on the basis of the budget passed by the Saxon state parliament (to M.F.), Deutsche Forschungsgemeinschaft (FE 1236/5-1, FE 1236/8-1 to M.F.), and Bundesministerium für Bildung und Forschung (01EJ2205A, FERROPath to M.F.).

## AUTHOR CONTRIBUTIONS

M.L. and J.A.O. conceived of the project, designed the experiments, and wrote the majority of the manuscript. All authors read, edited, and contributed to the manuscript. M.L. performed the majority of the experiments. C.D. purified FSP1 proteins used in biochemical assays, B.S. developed experiments in liposomes, and J.H. assisted with incucyte assays. M.W. and M.F. assisted with oxidized lipidomics analyses. K.N., Y.O., and S.K. provided key lipid standards.

## METHODS

### LC-MS based Lipidomics and Epi-Lipidomics

Materials used for liquid chromatography/mass spectrometry were: water (Fisher Scientific, W6-4), acetonitrile (Fisher Scientific, A955-4), 2-propanol (Supelco, 1.02781.4000), ammonium formate (Sigma-Aldrich, 70221-25G-F) and formic acid (Fisher Scientific, A117-50). Solvents for lipid extraction were tert-butyl methyl ether (Sigma-Aldrich, 34875) and methanol (Fisher Scientific, A456) that were spiked with 0.1% (w/v) 2,6-Di-tert-butyl-4-methylphenol (Sigma-Aldrich, B1378). Lipid internal standard mixture was SPLASH ® LIPIDOMIX ® Mass Spec Standard (Avanti Research, 330707).

### Lipid Extraction (for lipidomics and epi-lipidomics)

Confluent culture plates (10 cm or 6 cm diameter) were washed once with PBS and scraped into 1 ml ice-cold PBS. Cells were pelleted at 3,000xg for 5 min at 4C, the supernatant was removed, and cell pellets were stored at -80C. Before extraction, cell pellets were thawed at room temperature for 3 min and suspended in 50 µl PBS. Internal standards (3 µl SPLASH® LIPIDOMIX® per sample) dissolved in methanol were added directly to each cell suspension. Lipids were extracted by adding 1250 µl tert-butyl methyl ether and 375 µL methanol. The mixture was incubated on an orbital mixer for 1 h at room temperature (revolver shaker, 32 rpm). To induce phase separation, 315 µL water was added, and the mixture was incubated on for 10 min at room temperature (revolver shaker, 32 rpm). Samples were centrifuged at 15,000 x g at room temperature for 3 min. The upper organic phase was collected and subsequently dried *in vacuo* (Eppendorf concentrator 5301).

### Liquid Chromatography

Dried lipid extracts were reconstituted in 150 µl chloroform/methanol (2:1, v/v) and 20 µl of each extract was aliquoted in HPLC vials containing glass inserts. Pooled quality control (pooled QC) samples were generated by mixing equal volumes of each lipid extract followed by aliquotation in 20 µl aliquots. Aliquoted extracts and pooled QCs were dried *in vacuo* (Eppendorf concentrator 5301) and redissolved in 20 µl isopropanol for injection.

Lipids were separated by reversed phase liquid chromatography on a Vanquish Core (Thermo Fisher Scientific, Bremen, Germany) equipped with an Accucore C30 column (150 x 2.1 mm; 2.6 µm, 150 Å, Thermo Fisher Scientific, Bremen, Germany). Lipids were separated by gradient elution with solvent A (acetonitrile/water, 1:1, v/v) and B (isopropanol/acetonitrile/water, 85:10:5, v/v) both containing 5 mM ammonium formate and 0.1% (v/v) formic acid. Separation was performed at 50°C with a flow rate of 0.3 mL/min using the following gradient: 0-15 min – 25 to 86 % B (curve 5), 15-21 min – 86 to 100 % B (curve 5), 21-32 min – 100 % B isocratic, 32-32.1 min – 100 to 25 % B (curve 5), followed by 6 min re-equilibration at 25 % B.

### Mass Spectrometry

Reversed phase liquid chromatography was coupled on-line to a Q Exactive Plus Hybrid Quadrupole Orbitrap mass spectrometer (Thermo Fisher Scientific, Bremen, Germany) equipped with a HESI probe. Mass spectra were acquired in positive and negative modes with the following ESI parameters: sheath gas – 40 L/min, auxiliary gas – 10 L/min, sweep gas – 1 L/min, spray voltage – (+)3.5 kV (positive ion mode); (-)3.2 kV (negative ion mode), capillary temperature – 250 °C, S-lens RF level – 35 and aux gas heater temperature – 370 °C.

Data acquisition for lipid identification was performed in pooled QC samples by acquiring data in data dependent acquisition mode (DDA). DDA parameters featured a survey scan resolution of 140,000 (at *m/z* 200), AGC target 1e6, Maximum injection time 100 ms in a scan range of *m/z* 240-1200. Data dependent MS/MS scans were acquired with a resolution of 17,500, AGC target 1e5, Maximum injection time 60 ms, isolation window 1.2 *m/z* and stepped normalized collision energies of 10, 20 and 30. A data dependent MS2 was triggered (loop count 15) when an AGC target of 2e2 was reached followed by a Dynamic Exclusion for 10 s. All isotopes and charge states > 1 were excluded. All data was acquired in profile mode.

For deep lipidome profiling, iterative exclusion was performed using the IE omics R package. ^61^ This package generates a list for already fragmented precursors from a prior DDA run to be excluded from subsequent DDA runs ensuring a higher number of unique MS/MS spectra for deep lipidome profiling. After the initial DDA analysis of a pooled QC sample, the pooled QC was measured two more times but excluding all previously fragmentated precursor ions. Samples were analyzed both in positive and negative ion modes. Parameters for generating exclusion lists from previous runs were – RT window = 0.3; noiseCount = 15; MZWindow = 0.02 and MaxRT = 36 min.

Data for lipid quantification in individual samples was acquired in *Full MS* mode with following parameters – scan resolution of 140,000 (at *m/z* 200), AGC target 1e6, Maximum injection time 100 ms in a scan range of *m/z* 240-1200.

### Lipid Identification and Quantification

Lipostar (version 2.0, Molecular Discovery, Hertfordshire, UK) equipped with an *in house generated* structure database built in LipoStarDB manager was used. This database features fatty acids with no information on double bond regio- or stereoisomerism and covers glycerolipid, glycerophospholipid, sphingolipid and sterol ester lipid classes. The raw files were imported directly with a *Sample MS Signal Filter Signal Threshold* = 1000 for MS and a *Sample MS/MS Signal Filter Signal Threshold* = 10. Automatic peak picking was performed with an *m/z tolerance* = 5 ppm, *chromatography filtering threshold* = 0.97, *MS filtering threshold* = 0.97, *Signal filtering threshold* = 0. Peaks smoothing was performed using the *Savitzky-Golay* smoothing algorithm with a *window size* = 3, *degree* = 2 and *multi-pass iterations* = 3. Isotopes were clustered using a *m/z tolerance* = 5 ppm, *RT tolerance =* 0.25 min, *abundance Dev* = 40%, *max charge* = 1. Peak alignment between samples using an *m/z tolerance* = 5 ppm and an *RT tolerance* = 0.25 min. A gap filler with an *RT tolerance* = 0.05 min and a *signal filtering threshold* = 0 with an anti-Spike filter was applied.

For lipid identification, a “MS/MS only” filter was applied to keep only features with MS/MS spectra for identification. Triacylgylcerols, diacylglycerols and steryl esters were identified as [M+NH4]+ adducts. Lysophosphatidylcholines; lysophosphatidylethanolamines; Acyl, ether- and vinyl ether-PE, phosphatidylserines, phosphatidylinositols, ceramides and sphingomyelins were analyzed as [M+H]+ adducts. Acyl-, ether- and vinyl ether-Phosphatidylcholines were identified as [M-H]- adducts.^62^ Following parameters were used for lipid identification: 5 ppm precursor ion mass tolerance and 20 ppm product ion mass tolerance. Automatic approval was performed to keep structures with quality of 3-4 stars. Identifications were refined using manual curation and Kendrick mass defect analysis and lipids that were not following linear Kendrick mass defect-retention time correlations were excluded as false positives.

Quantification was performed by peak integration of the extracted ion chromatograms of single lipid adducts. Peak integration was manually curated and adjusted. Identified lipids were normalized to peak areas of added internal standards to decrease analytical variation and eventually normalized to protein concentrations of cell pellets after lipid extraction.

### Statistical analysis of lipidomics data

Metaboanalyst 5.0 (https://new.metaboanalyst.ca/home.xhtml) was used for statistical analysis and data transformation. Raw lipidomics data was imported into the “Statistical Analysis [one factor]” analysis pipeline. Data was transformed by “Log transformation (base 10)” and scaled by “Auto scaling” for statistical analysis. The transformed data and calculated statistics were exported as .csv files and plotted in GraphPad Prism 10.2.2 (GraphPad Software).

### Comprehensive Epi-Lipidomics (oxidized lipid analysis) for analysis of FSP1 mutants

#### Prediction of sample specific oxidized lipidome data-dependent acquisition

For the semi-targeted identification of oxidized lipids from cell extracts we followed a protocol as published before.^58^ To narrow down the number of potential lipid peroxidation products, we chose a subset of lipids for *in silico* oxidation. This lipid subset consisted of the top 10 lipids with the highest intensity of PC, P-PC, O-PC, PE, P-PE, O-PE and the top 20 lipids with the highest intensity of TG and CE lipid classes identified in each FSP1 mutant cell lipidome. Additionally, we incorporated the top 10 lipids whose intensity correlates positively with sensitivity to cell death induced by RSL3 in each FSP1 mutant cell line. This led to a target list for *in silico* oxidation of 176 lipids in total represented by 77 phospholipids and 99 non-polar lipids.

*In silico* oxidation on this subset of rationally selected lipids was performed with LPPTiger2^63,64^ utilizing following parameters: “Max modification site = 2”, “Max total O = 3”, “max OH = 2”, “max keto = 1”, “max OOH = 1”, max epoxy = 0”. The predicted epilpidome was refined by the unmodified lipid list generated in the step described above. An inclusion list was generated featuring the [M+Na]+ adducts of these entries as the generated fragments of sodiated oxidized lipid ions are most informative for oxidized lipid identification.

#### Liquid Chromatography

Pooled QC samples were dissolved in 20 µl isopropanol for injection.

Lipids were separated by reversed phase liquid chromatography on a Vanquish Horizon (Thermo Fisher Scientific, Bremen, Germany) equipped with an Accucore C30 column (150 x 2.1 mm; 2.6 µm, 150 Å, Thermo Fisher Scientific, Bremen, Germany). Lipids were separated by gradient elution with solvent A (acetonitrile/water, 1:1, v/v) and B (isopropanol/acetonitrile/water, 85:10:5, v/v) both containing 5 mM ammonium formate and 0.1% (v/v) formic acid. Separation was performed at 50°C with a flow rate of 0.3 mL/min using the following gradient: 0-20 min – 10 to 80 % B (curve 5), 20-24 min – 80 to 95 % B (curve 5), 24-27 min – 95 to 100 % B (curve 5), 27 min – 32 min 100 % B isocratic, 32-32.1 min – 100 to 10 % B (curve 5), followed by 7.9 min re-equilibration at 10 % B.

#### Data dependent acquisition settings to target in silico oxidized epilipidome

Reversed phase liquid chromatography was coupled on-line to an Orbitrap Exploris 240 Hybrid Quadrupole Orbitrap mass spectrometer (Thermo Fisher Scientific, Bremen, Germany) equipped with a HESI probe. Mass spectra were acquired in positive and negative modes with the following ESI parameters: sheath gas – 40 arb, auxiliary gas – 10 arb, sweep gas – 1 arb, spray voltage – (+)3.5 kV (positive ion mode); (-)2.5 kV (negative ion mode), ion transfer tube temperature – 300 °C, S-lens RF level – 35 and vaporizer temperature – 370 °C, EASY-IC was set to run start. Data acquisition for oxidized lipid identification was performed in quality control samples by acquiring data in data dependent acquisition mode (DDA) with an inclusion list of predicted oxidized lipids. DDA parameters featured a survey scan resolution of 60,000 (at *m/z* 200), AGC target 1e6, Maximum injection time auto in a scan range of *m/z* 500 - 980 in negative ion mode (0 - 22 min) and *m/z* 480 – 1200 in positive ion mode (22 - 40 min). Data dependent MS/MS scans were acquired with a resolution of 60,000, normalized AGC target 100%, maximum injection time 200 ms, number of dependent scans 6, isolation window 1.5 *m/z*, first fixed mass *m/z* 100, microscans 2 and normalized HCD collision energies of 22,32,43% in negative ion mode and32,43,54 % in positive ion mode. Dynamic Exclusion was set to exclude after 5 times withing 6s for 5s with a mass tolerance of ±5ppm. All isotopes and charge states > 1 were excluded. All data was acquired in profile mode.

#### Identification of oxidized lipids from data dependent acquisition dataset

MSMS scans containing informative fragments were extracted using LPPtiger^63^ in lipid identification mode. The .raw files were converted to .mzML using MSConvertGUI from ProteoWizard 3.0.9134.^65^ mzML files were uploaded to LPPTiger2 and MSMS spectra containing oxidized lipid candidates were identified with following settings: “MS1 tolerance = 5 ppm”, “MSMS tolerance = 20 ppm”, “Precursor intensity = 100”, “Precursor mz window = *m/*z 480-1100”, “Selection window = *m/z* 1.2” with a “Score filter > 60”, “Isotope Score > 80”, “Rank Score > 40” and considering peaks with intensity > 1%.

To remove false-positive MSMS candidates and improve identification confidence, the spectra were filtered manually and MSMS candidates that did not contain an “oxidized fatty acid specific fragment” were removed. For oxTG and oxCE,n “oxidized fatty acid specific fragment” was defined as a [M+Na]+ or [M+H]+ adduct of an intact oxidized fatty acid (e.g. [FA18:2<OOH>+Na]+) or an oxidized fatty acid fragment that after fragmentation retained at least one oxygen atom (e.g. [FA18:2<OOH>-H2O+H]+). MSMS candidates containing fragments originating from oxidized fatty acids that do not retain additional oxygen atoms after fragmentation were removed (e.g. acid (e.g. [FA18:2<OH>-H2O+H]+) as these spectra cannot be used to confidently determine the presence of a fatty acid specific modification. For oxPC and oxPE an “oxidized fatty acid fragment” was defined as a [M-H]-adduct of an intact oxidized fatty acid (e.g. [FA18:2<OOH>-H]-) or an oxidized fatty acid fragment that after fragmentation retained at least one oxygen atom (e.g. [FA18:2<OOH>-H2O-H]-). MSMS candidates containing fragments originating from oxidized fatty acids that do not retain additional oxygen atoms after fragmentation were removed (e.g. acid (e.g. [FA18:2<OH>-H2O-H]-) as these spectra cannot be used to confidently determine the presence of a fatty acid specific modification. Isomeric species were removed to result in a final list covering 82 *m/z* values representing oxidized lipids.

#### Relative quantification of oxidized lipids by parallel reaction monitoring

Lipids were dissolved in isopropanol and separated and ionized as described above for data dependent acquisition.

Optimized data acquisition settings for parallel reaction monitoring were: “Isolation window = 1.5 *m/z*”, “Resolution = 15,000”, “maximum IT = 100 ms”, “microscans = 1” featuring a single collision energy. Transitions were quantified with Skyline^60^ utilizing a 5 ppm mass accuracy on a centroided fragment signal. Confidence of quantification was improved by observing transitions of [M+Na]+ → [oxFA+Na]+ or [oxFA+H]+ depending on the targeted lipid.

Signal intensities of each oxidized lipid were normalized to isotopically labeled standards of their respective lipid class in the SPLASH LIPIDOMIX (Avanti Research, 330707) and finally normalized to the protein concentration in each cell pellet previously determined by a bicinchoninic acid protein assay kit (Thermo Fisher Scientific, PI23227).

#### Statistical analysis of Epi-lipidomics data

Metaboanalyst 5.0 (https://new.metaboanalyst.ca/home.xhtml) was used for statistical analysis and data transformation. Raw lipidomics data was imported into the “Statistical Analysis [one factor]” analysis pipeline. Data was transformed by “Log transformation (base 10)” and scaled by “Auto scaling” for statistical analysis. The transformed data and calculated statistics were exported as .csv files and plotted in GraphPad Prism 10.2.2 (GraphPad Software).

### Epi-Lipidomics of oxidized non-polar lipids in PUFA induced LD damage model

#### Prediction of sample specific oxidized lipidome data-dependent acquisition

For the untargeted identification of oxidized lipids from cell extracts we followed a protocol as published before.^60^ To narrow down the number of potential lipid peroxidation products, we chose a subset of lipids for in silico oxidation. This lipid subset consisted of the top 15 lipids with the highest intensity of TG and CE lipid classes in either U-2 OS wild type cells treated with 200 µM arachidonic acid or U-2 OS TetR and U-2 OS FSP1 KO cells treated with 20 µM arachidonic acid. Additionally, we incorporated the lipids that were significantly downregulated (i.e. log2 fold change > 2, *p* < 0.05) by FSP1 inhibition via FSEN1 treatment in the arachidonic acid treated conditions. This lead to a target list for in silico oxidation of 95 lipids in total represented by 18 CE and 77 TG lipids.

*In silico* oxidation on this subset of rationally selected lipids was performed with LPPTiger^63^ utilizing following parameters: “Max modification site = 2”, “Max total O = 3”, “max OH = 2”, “max keto = 1”, “max OOH = 1”, max epoxy = 0”. The predicted epilpidome was refined by the unmodified lipid list generated in the step described above. An inclusion list was generated featuring the [M+Na]+ adducts of these entries as the generated fragments of sodiated oxidized lipid ions are most informative for oxidized lipid identification.

#### Liquid Chromatography

Dried lipid extracts were reconstituted in 60 µl chloroform/methanol (2:1, v/v) and 10 µl of each extract was aliquoted in HPLC vials containing glass inserts. Quality control samples were generated by mixing equal volumes of each lipid extract followed by aliquotation in 10 µl aliquots. Aliquoted extracts were dried *in vacuo* (Eppendorf concentrator 5301) and redissolved in 20 µl isopropanol for injection and 5 µl were loaded onto the column.

Lipids were separated by reversed phase liquid chromatography on a Vanquish Core (Thermo Fisher Scientific, Bremen, Germany) equipped with an Accucore C30 column (150 x 2.1 mm; 2.6 µm, 150 Å, Thermo Fisher Scientific, Bremen, Germany). Lipids were separated by gradient elution with solvent A (acetonitrile/water, 1:1, v/v) and B (isopropanol/acetonitrile/water, 85:10:5, v/v) both containing 5 mM ammonium formate and 0.1% (v/v) formic acid. Separation was performed at 50°C with a flow rate of 0.3 mL/min using the following gradient: 0-15 min – 25 to 86 % B (curve 5), 15-21 min – 86 to 100 % B (curve 5), 21-32 min – 100 % B isocratic, 32-32.1 min – 100 to 25 % B (curve 5), followed by 6 min re-equilibration at 25 % B.

#### Data dependent acquisition settings to target in silico oxidized epilipidome

Reversed phase liquid chromatography was coupled on-line to an QExactive Plus Hybrid Quadrupole Orbitrap mass spectrometer (Thermo Fisher Scientific, Bremen, Germany) equipped with a HESI probe. Mass spectra were acquired in positive and negative modes with the following ESI parameters: sheath gas – 40 L/min, auxiliary gas – 10 L/min, sweep gas – 1 L/min, spray voltage – (+)3.5 kV (positive ion mode, capillary temperature – 250 °C, S-lens RF level – 35 and aux gas heater temperature – 370 °C.

Data acquisition for oxidized lipid identification was performed in quality control samples by acquiring data in data dependent acquisition mode (DDA). DDA parameters featured a survey scan resolution of 140,000 (at m/z 200), AGC target 1e6 Maximum injection time 100 ms in a scan range of m/z 500-1200. Data dependent MSMS scans were acquired with a resolution of 17,500, AGC target 1e6, Maximum injection time 200 ms, loop count 7, isolation window 1.2 m/z, fixed fist mass = 100 and normalized collision energy of 30. A data dependent MSMS was triggered when an AGC target of 1e1 was reached followed by a Dynamic Exclusion for 3 s. All isotopes and charge states > 1 were excluded. All data was acquired in profile mode.

#### Identification of oxidized lipids from data dependent acquisition dataset

MSMS scans containing informative fragments were extracted using LPPtiger^63^ in lipid identification mode. The .raw files were converted to .mzML using MSConvertGUI from ProteoWizard 3.0.9134^65^. mzML files were uploaded to LPPTiger and MSMS spectra containing oxidized lipid candidates were identified with following settings: “MS1 tolerance = 5 ppm”, “MSMS tolerance = 20 ppm”, “Precursor intensity = 100”, “Precursor mz window = *m/*z 500-1100”, “Selection window = *m/z* 1.2” with a “Score filter > 60”, “Isotope Score > 80”, “Rank Score > 40” and considering peaks with intensity > 1%.

To remove false-positive MSMS candidates and improve identification confidence, the spectra were filtered manually and MSMS candidates that did not contain an “oxidized fatty acid specific fragment” were removed. An “oxidized fatty acid specific fragment” was defined as a [M+Na]+ or [M+H]+ adduct of an intact oxidized fatty acid (e.g. [FA18:2<OOH>+Na]+) or an oxidized fatty acid fragment that at least retained one oxygen atom after fragmentation acid (e.g. [FA18:2<OOH>-H2O+H]+). MSMS candidates containing fragments originating from oxidized fatty acids that do not retain additional oxygen atoms after fragmentation were removed (e.g. acid (e.g. [FA18:2<OH>-H2O+H]+) as these spectra cannot be used to confidently determine the presence of a fatty acid specific modification. All entries were sorted and isomeric species were removed to result in a final inclusion list covering 16 *m/z* values representing oxidized lipids.

#### Relative quantification of oxidized lipids by parallel reaction monitoring

Lipids were dissolved in isopropanol and separated and ionized as described above for data dependent acquisition.

Confidence of quantification was improved by observing transitions of [M+Na]+ → [oxFA+Na]+ or [oxFA+H]+ depending on the targeted lipid.

Optimized data acquisition settings for parallel reaction monitoring were: “Isolation width = 1.2 *m/z*”, “Resolution = 17,500”, “maximum IT = 200 ms”, “microscans = 1”, “maximum IT = 200 ms” featuring a “normalized collision energy = 30”. Transitions were quantified with Skyline^66^ utilizing a 5 ppm mass accuracy on a centroided fragment signal. Signal intensities of each oxidized lipid were normalized to isotopically labeled standards of their respective lipid class in the SPLASH LIPIDOMIX (Avanti Research, 330707).

#### Statistical analysis of Epi-lipidomics data

Metaboanalyst 5.0 (https://new.metaboanalyst.ca/home.xhtml) was used for statistical analysis and data transformation. Raw lipidomics data was imported into the “Statistical Analysis [one factor]” analysis pipeline. Data was transformed by “Log transformation (base 10)” and scaled by “Auto scaling” for statistical analysis. The transformed data and calculated statistics were exported as .csv files and plotted in GraphPad Prism 10.2.2 (GraphPad Software).

### Relative quantification of redox active lipids

Wild type U-2 OS cells were treated with 200 uM oleate (Nu-Chek Prep) and 0.1% fatty acid free BSA (Sigma Aldrich, A3803). Lipid droplets were extracted as described below in method “Lipid droplet enrichment/flotation”.

Lipophilic radical trapping antioxidants were extracted by addition of 600 µl methanol (Fisher Scientific, A456) + 0.1% (v/v) hydrochloric acid (Fisher Scientific, A144) and 600 µl hexane (Fisher Scientific, H302) to a 600 µl LD fraction. Samples were vortexed vigorously for 60s at room temperature and centrifuged at 2,000xg for 3 min at 4C. The upper hexane fraction was collected and transferred to an HPLC glass vial. The upper hexane phase was dried *in vacuo* (vacuum concentrator 5301, Eppendorf). For LCMS analysis, the dried fraction was dissolved in 50 µl isopropanol (Supelco, 1.02781.4000) and 10 µl were injected for analysis. Coenzyme Q10, Q9, Q8, Vitamin K1 (phylloquinone), Vitamin K2 (menaquinone 4), (+)-α-tocopherol were analyzed via parallel reaction monitoring.

Lipids were separated by reversed phase liquid chromatography on a Vanquish Core (Thermo Fisher Scientific, Bremen, Germany) equipped with an Accucore C30 column (150 x 2.1 mm; 2.6 µm, 150 Å, Thermo Fisher Scientific, Bremen, Germany). Lipids were separated by gradient elution with solvent A (acetonitrile/water, 1:1, v/v) and B (isopropanol/acetonitrile/water, 85:10:5, v/v) both containing 5 mM ammonium formate and 0.1% (v/v) formic acid. Separation was performed at 50°C with a flow rate of 0.3 mL/min using the following gradient: 0-15 min – 25 to 86 % B (curve 5), 15-21 min – 86 to 100 % B (curve 5), 21-32 min – 100 % B isocratic, 32-32.1 min – 100 to 25 % B (curve 5), followed by 6 min re-equilibration at 25 % B.

Reversed phase liquid chromatography was coupled on-line to a Q Exactive Plus Hybrid Quadrupole Orbitrap mass spectrometer (Thermo Fisher Scientific, Bremen, Germany) equipped with a HESI probe. Mass spectra were acquired in positive and negative modes with the following ESI parameters: sheath gas – 40 L/min, auxiliary gas – 10 L/min, sweep gas – 1 L/min, spray voltage – (+)3.5 kV (positive ion mode); (-)3.2 kV (negative ion mode), capillary temperature – 250 °C, S-lens RF level – 35 and aux gas heater temperature – 370 °C.

Lipids were analyzed by parallel reaction monitoring and individual ions were isolated with an isolation window of 1.2 m/z and MSMS spectra were acquired with a AGC target of 1e6 with a maxIT of 100 ms and a resolution of 17,500. Quantification was of each lipid was optimized and final settings were:

| Analyte | tR<br>[min] | collision<br>energy<br>[a.u.] | Precursor<br>Formula | Adduct | Product<br>Formula | Product<br>m/z | Adduct |
| --- | --- | --- | --- | --- | --- | --- | --- |
| CoQ10 | 22.4 | 15 | C59H90O4 | [M+NH4]<sup>+</sup> | C10H12O4 | 197.0808 | [M+H]<sup>+</sup> |
| CoQ10H2 | 22.1 | 15 | C59H92O4 | [M+NH4]<sup>+</sup> | C10H12O4 | 197.0808 | [M+H]<sup>+</sup> |
| CoQ9 | 21.4 | 15 | C54H82O4 | [M+NH4]<sup>+</sup> | C10H12O4 | 197.0808 | [M+H]<sup>+</sup> |
| CoQ9H2 | 21.1 | 15 | C54H84O4 | [M+NH4]<sup>+</sup> | C10H12O4 | 197.0808 | [M+H]<sup>+</sup> |
| CoQ8 | 20.2 | 15 | C49H74O4 | [M+NH4]<sup>+</sup> | C10H12O4 | 197.0808 | [M+H]<sup>+</sup> |
| Vitamin K1 | 17.5 | 25 | C31H46O2 | [M+H]<sup>+</sup> | C12H10O2 | 187.0754 | [M+H]<sup>+</sup> |
| Menaquinone 4 | 14.8 | 25 | C31H40O2 | [M+H]<sup>+</sup> | C12H10O2 | 187.0754 | [M+H]<sup>+</sup> |
| α-tocopherol | 13.2 | 20 | C29H50O2 | [M+H]<sup>+</sup> | C10H12O2 | 165.091 | [M+H]<sup>+</sup> |

### Cellular and *in vitro* assays

#### Cell culture

U-2OS cells parental cells were obtained from the UC Berkeley Cell Culture Facility. Cells were cultured in DMEM with l-glutamine and without sodium pyruvate (Corning, Cat# 10-017-CM) containing 10% fetal bovine serum (GemCell, 100-500). All cells were maintained at 37C and 5% CO2.

FSP1 mutant expression was induced in U-2 OS cells with 10 ng/ml doxocycline (Sigma Aldrich, D9891) for 48 hours prior to the respective experiment.

#### Immunodetection (Western Blotting) of FSP1

Cells were cultured in 10 cm culture dishes. Before collection, cells were washed with 5 ml PBS and scraped into 1 ml PBS. Cells were pelleted by centrifugation at 3,000xg for 3 min at 4C and the supernatant was discarded. For lysis, cells were resuspended in 100 µl RIPA buffer (Pierce, 89901) with 1x protease inhibitor cocktail (Pierce, A32955) and sonicated by probe sonication for 30 seconds with 15% amplitude on a Branson 150 sonicator equipped with a sonication probe (Branson Ultrasonics, 4C15). Protein concentration was determined via a bicinchoninic acid protein assay kit (Thermo Fisher Scientific, PI23227).

For gel electrophoresis, cell lysates were diluted with 1% SDS and 10-15 µg of protein were loaded on a 4-20% gradient gel (Bio-Rad, 4561094) and separated at 90V for 10 min followed by 250V for 20 min. Proteins were transferred to nitrocellulose membranes (Bio-Rad, 1704158) using the TransblotTurbo system (Bio-Rad) at mixed molecular weight settings. Membranes were blocked with 5% milk in PBS containing 0.1% Tween-20 (Sigma Aldrich, P7949) for 1 hour at room temperature. Blots were washed three times with PBS containing 0.1% Tween-20. Primary antibodies were diluted 1:1,000 in 5% BSA (Fisher Scientific, BP9703) in PBS containing 0.1% Tween-20 and blots were incubated with primary antibodies overnight at 4C. Next, blots were washed three times with PBS containing 0.1% Tween-20. Secondary antibodies were diluted 1:25,000 in 5% BSA in PBS containing 0.1% Tween-20. After washing three times with PBS containing 0.1% Tween-20, blots were imaged on a Li-Cor imaging system (Li-Cor, Odyssey).

#### Cell death analysis

Cells were seeded into 96 well plates (Corning, 3904) at 15% confluency in 10 ng/ml doxycycline hyclate (Sigma Aldrich, D9891) for 48 hours for induced FSP1 expression or 30% confluency for 24 hours in wild type cells prior to the respective experiment. Lipid droplets were induced by treating cells with 200 uM of the respective fatty acid (Nu-Chek Prep) and 0.1% fatty acid free BSA (Sigma Aldrich, A3803).

Ferroptosis initiators were dissolved in DMSO and mixed with the cell death marker sytox green (Thermo Fisher, S34860) in standard culture medium to yield a 2x stock solution. 100 µl of the 2x stock solution were added on top of 100 µl media in each well. Cell death propagation was followed by time resolved fluroescence microscopy using an IncuCyte S3 imaging system (Essen Bioscience).

Fatty acids were dissolved in ethanol and mixed with the cell death marker sytox green (Thermo Fisher, S34860) in standard culture medium to yield a 2x stock solution. To analyze influence of lipid metabolism regulators, cells were co-treated with 5 µM FSEN1 (Cayman Chemicals, 38025), 1 µg/ml Triacsin C (EnzoLifeSciences, BML-EI218), 10 uM NG-497 (Cayman Chemicals, 36886), 20 µM T-863 (Cayman Chemicals, 25807), 15 µM A-922500 (Cayman Chemicals, 10012708) or 10 µM PF-06424439 (Cayman Chemicals, 17680).

#### Lipid droplet imaging and quantification

Cells were seeded into 96 well plates (Corning, 3904) at 30% confluency for 24 hours in wild type cells prior to imaging. Lipid droplets were induced by treating cells with 200 uM of the respective fatty acid (Nu-Chek Prep) and 0.1% fatty acid free BSA (Sigma Aldrich, A3803). To block TG synthesis, DGAT inhibitors were added at the same time as fatty acids. DGAT1 was inhibited by 15 µM A-922500 (Cayman Chemicals, 10012708) and DGAT2 was inhibited by 10 µM PF-06424439 (Cayman Chemicals, 17680).

For lipid droplet imaging, cells were stained with 1 ug/ml BODIPY 493/503 (Invitrogen, D3922) for 25 min and Hoechst (Invitrogen, H3570) following vendor’s recommendation. Media was removed and replaced with pre-warmed phenol red free medium (Cytiva, SH30284.02) supplemented with 10% fetal bovine serum (Gemini, 100-500).

BODIPY 493/503 was imaged on an Opera Phenix (Revvity) with laser excitation at 488 nm and emission was read at 500-550 nm. Hoechst was imaged using the standard DAPI filter sets.

Lipid droplets number and area was quantified using Harmony data analysis software (Revvity).

#### Lipid droplet enrichment/flotation

1.25 million cells were seeded in a p150 cell culture dish and grown for 24 hours. Lipid droplet formation was induced with 200 uM oleate (NuChek-Prep, U-46-A) and 0.1% fatty acid free BSA (Sigma-Aldrich, A3803) for 24 hours. Cells were collected by scraping into 3 ml PBS and pelleted by centrifugation at 1,000xg for 15 min at 4C. Cell pellets were resuspended in 2 ml hypotonic lysis medium (20 mM Tris-HCl pH 7.4, 1 mM EDTA) with protease inhibitor cocktail (Thermo Fisher, A32955) and incubate for 10 min on ice. Cells were lysed by pushing cell suspension through a 23G needle (BD PrecisionGlide, 305145) for a total of 8 times. Centrifuge cell lysate at 1,000xg for 10 min at 4C. For lipid droplet flotation the supernatant was transferred to a 13.2 ml centrifuge tube (Beckman Coulter, 344059) and mixed with 60% sucrose (Fisher Scientific, S5-500) in hypotonic lysis medium to achieve a final concentration of 20% sucrose. This solution was overlayed with an ice-cold solution of 5% sucrose in hypotonic lysis medium followed by an ice-cold solution of hypotonic lysis medium without sucrose. Sucrose gradients were centrifuged at 28,000xg for 30 min at 4C in an SW-41TI rotor (Beckman Coulter). Collect the lipid droplet containing buoyant fraction was collected as described previously^67^ and stored at -80C until analysis.

#### Absolute quantification of coenzyme q10 in lipid droplets

600 µl lipid droplet fraction was dried at room temperature for 5 min and 2.5 pmol CoQ10-d6 (Cayman Chemicals, 30958) in isopropanol (Supelco, 1.02781.4000) was added and vortexed. Coenzyme Q10 was extracted as described before^68^ by addition of 600 µl methanol (Fisher Scientific, A456) + 0.1% (v/v) hydrochloric acid (Fisher Scientific, A144) and 600 µl hexane (Fisher Scientific, H302). Samples were vortexed vigorously for 60s at room temperature and centrifuged at 2,000xg for 3 min at 4C. The upper hexane fraction was collected and transferred to an HPLC glass vial. The upper hexane phase was dried *in vacuo* (vacuum concentrator 5301, Eppendorf). For LCMS analysis, the dried fraction was dissolved in 50 µl isopropanol (Supelco, 1.02781.4000) and 10 µl were injected for analysis.

Lipids were separated by reversed phase liquid chromatography on a Vanquish Core (Thermo Fisher Scientific, Bremen, Germany) equipped with an Accucore C30 column (150 x 2.1 mm; 2.6 µm, 150 Å, Thermo Fisher Scientific, Bremen, Germany). Lipids were separated by gradient elution with solvent A (acetonitrile/water, 1:1, v/v) and B (isopropanol/acetonitrile/water, 85:10:5, v/v) both containing 5 mM ammonium formate and 0.1% (v/v) formic acid. Separation was performed at 50°C with a flow rate of 0.3 mL/min using the following gradient: 0-15 min – 25 to 86 % B (curve 5), 15-21 min – 86 to 100 % B (curve 5), 21-32 min – 100 % B isocratic, 32-32.1 min – 100 to 25 % B (curve 5), followed by 6 min re-equilibration at 25 % B.

Reversed phase liquid chromatography was coupled on-line to a Q Exactive Plus Hybrid Quadrupole Orbitrap mass spectrometer (Thermo Fisher Scientific, Bremen, Germany) equipped with a HESI probe. Mass spectra were acquired in positive and negative modes with the following ESI parameters: sheath gas – 40 L/min, auxiliary gas – 10 L/min, sweep gas – 1 L/min, spray voltage – (+)3.5 kV (positive ion mode); (-)3.2 kV (negative ion mode), capillary temperature – 250 °C, S-lens RF level – 35 and aux gas heater temperature – 370 °C.

| Analyte | tR<br>[min] | collision<br>energy [a.u.] | Precursor<br>Formula | adduct | Product<br>Formula | Product<br>m/z | adduct |
| --- | --- | --- | --- | --- | --- | --- | --- |
| CoQ10 | 22.4 | 15 | C59H90O4 | [M+NH4]<sup>+</sup> | C10H12O4 | 197.0808 | [M+H]<sup>+</sup> |
| CoQ10H2 | 22.1 | 15 | C59H92O4 | [M+NH4]<sup>+</sup> | C10H12O4 | 197.0808 | [M+H]<sup>+</sup> |
| CoQ10-d6 | 22.4 | 15 | C59H84D6O4 | [M+NH4]<sup>+</sup> | C10H6D6O4 | 203.11904 | [M+H]<sup>+</sup> |

For absolute quantification, peak areas of CoQ10 and CoQ10H2 were divided by CoQ10-d6 and multiplied with the known concentration of CoQ10-d6.

#### Thin layer chromatography

Lipids were separated via thin layer chromatography on HPTLC Silica gel 60 plates with a preconcentration zone (Supelco, 1.13748.0001). Lipids were dissolved in chloroform/methanol (2:1, v/v) and applied onto the TLC with glass capillaries (CAMAG, 022.7729). Polar lipids were separated with triethylamine/chloroform/ethanol/water (5:5:5:1, v/v) and non-polar lipids were separated with hexane/diethyl ether/acetic acid (8:2:1, v/v) in a twin trough chamber (CAMAG, 022.5155) that was equilibrated with the eluent for 10 min prior to separation.

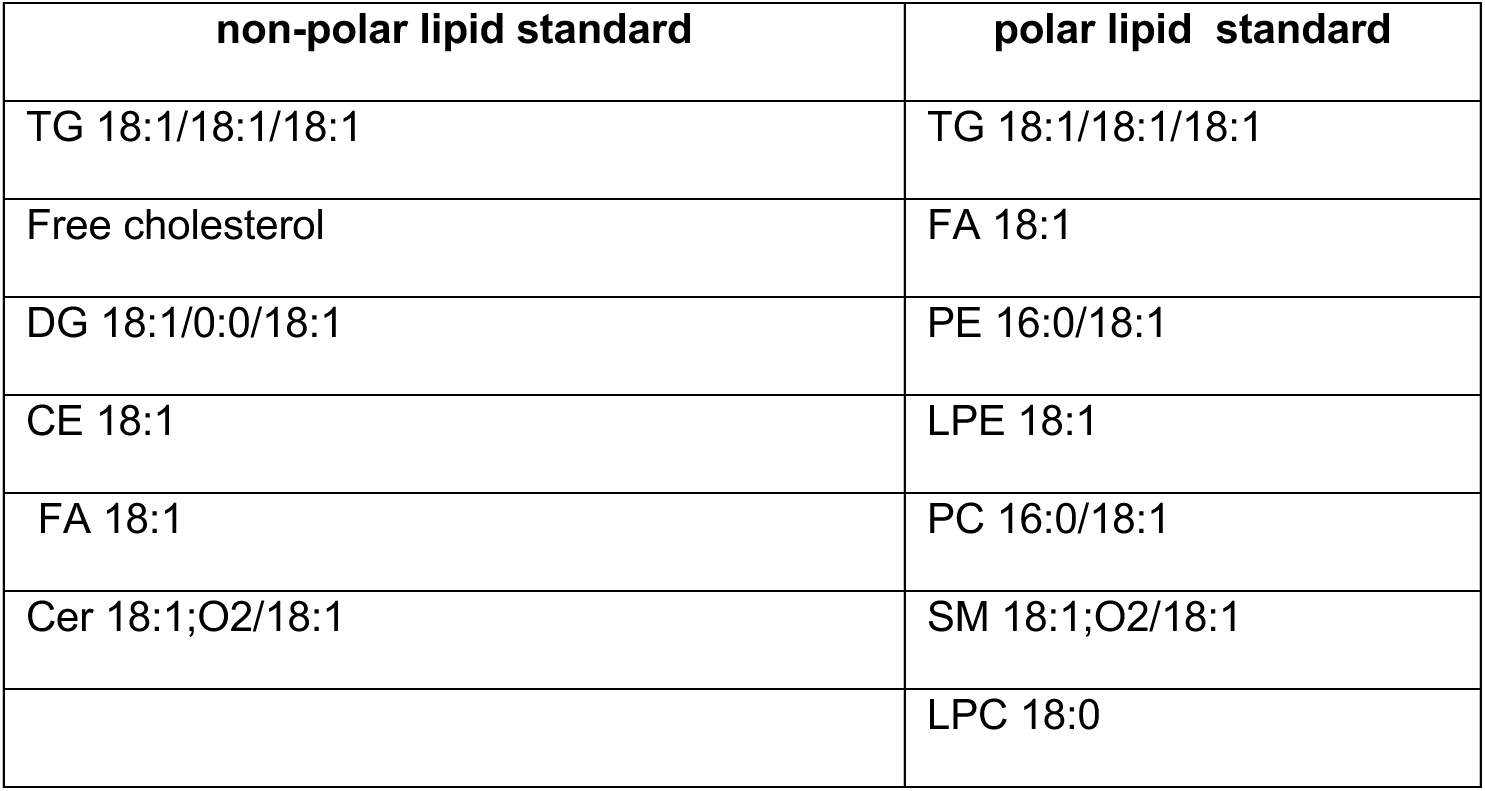

After separation plates were dried on air for 10 min and lipids were visualized by dipping plates for 2 seconds into a solution of primuline (Sigma Aldrich, 206865) (0.05%, w/v) in acetone/water (8:2, v/v). Plates were dried on air for 45 min and imaged on a ChemiDoc MP (BioRad) using blue epi light illumination and an emission filter of 530/28. Densitometric analysis was performed with Image Lab (Version 5.2.1, Bio-Rad).

#### Solid phase extraction for pre-analytical lipid separation

Lipids were separated into a fatty acid, polar lipid, and non-polar lipid fractions via solid phase extraction on an aminopropyl STRATA NH2 column (500 mg, 55 um, 70A, 8B-S009-HBJ, Phenomenex) as described previously.^69^ Columns were equilibrated with 4 ml hexane lipid extracts were loaded onto the column in 400 ul choloroform/methanol (2:1, v/v). Non-polar lipids were eluted with choloroform/isopropanol (2:1, v/v). Fatty acids were eluted with 8 ml diethyl ether/acetic acid (100:2, v/v) and polar lipids were eluted with 4 ml methanol. The collected fractions were dried *in vacuo* (Eppendorf concentrator 5301).

#### Gas-chromatography-flame ionization detection for absolute fatty acid quantification

Fatty acid composition of polar lipid and non-polar lipid fractions was analyzed at OmegaQuant (Sioux Falls, South Dakota) via gas chromatography coupled to flame ionization detection. Esterified fatty acids were hydrolyzed and methylated with an internal protocol and fatty acid methyl esters were separated on a GC2010 gas chromatograph (Shimadzu) equipped with a SP2560 fused silica capillary column (100-m, 0.25 mm internal diameter, 0.2 um film thickness, Supelco). Fatty acids were identified based on characteristic elution times of a fatty acid standard mixture (GLC OQ-A, Nu-Check Prep) and quantified based on external calibration.

#### Recombinant FSP1 production

Expression vectors were transformed into LOBSTR-BL21 (DE3) cells (Kerafast, EC1002) and grown overnight in LB broth at 37°C wile shaking (280 rpm). Cultures were diluted 1:100 into 1 L LB broth the next day and grown at 37°C until an OD600 of 0.6 (NanoDrop One, Thermo Fisher,13-400-518). Protein expression was induced with 0.7 mM IPTG, and cultures were incubated at 30°C with shaking (280 rpm) overnight. Cells were pelleted by centrifugation at 4500 rpm for 45 min at 4°C and resuspended in 25 mL lysis buffer (50 mM KH2PO4, 300 mM KCl, 10% glycerol, 30 mM imidazole, 1 mg/mL lysozyme, 1 mM PMSF, pH 8.0). After 10 min incubation at 37°C with shaking (280 rpm), cells were lysed by passing through a microfluidizer (Microfluidics LM10) 3× at 15,000 PSI and clarified by ultracentrifugation at 50,000 × g for 30 min at 4°C. Clarified lysate was loaded onto an Econo-Pac column (Bio-Rad, Cat# 732-1010) containing 1 mL HisPur Ni-NTA agarose resin (Thermo Fisher, 88221) equilibrated in wash buffer (50 mM KH2PO4, 300 mM KCl, 10% glycerol, 30 mM imidazole, pH 8.0). The column was washed 4× with 2 mL wash buffer, and proteins were eluted in 5 mL elution buffer (same as wash buffer but with 250 mM imidazole). Eluted proteins were buffer-exchanged into SEC buffer (50 mM HEPES, 100 mM KCl, 2 mM DTT, pH 8.0, spiked with 15% glycerol and 10 mM DTT) using an Amicon Ultra-15 filter (Sigma-Aldrich, UFC910008) and concentrated to 3 mL.

Proteins were purified via size exclusion chromatography on a HiLoad 16/600 Superdex 75 Pg column (Sigma-Aldrich, GE28-9893-33) in SEC buffer using a GE Akta Pure FPLC. Fractions (1.6 mL) were pooled based on SDS-PAGE purity (Coomassie-stained), concentrated to 2 mg/mL, and snap-frozen in 50 μL aliquots. Protein concentration was determined using a Pierce BCA protein assay kit (Thermo Fisher, PI23227).

#### FSP1 activity assay - NADH reducing activity

The ability of recombinant wild type FSP1 or the mutant FSP1(E156A) activity to consume NADH was tested in a 96 well format. Recombinant protein (25 nM) was mixed with the soluble CoQ10 derivative CoQ1 (Sigma Aldrich, C7956) in PBS. Immediately before the measurement, NADH (EMD Millipore, 481913) dissolved in PBS was added to result in a final concentration of 500 uM and reaction progress was followed using a TECAN Spark plate reader followed by measuring the absorbance of NADH at 340 nm every 15 seconds for 15 minutes. Data was plotted and analyzed with Graphpad Prism (10.2.2).

#### FSP1 activity assay (CoQ reduction activity)

The ability of recombinant wild type FSP1 or the mutant FSP1(E156A) activity to reduce oxidized CoQ10 was tested in a 96 well format. Recombinant protein (25 nM) was mixed with NADH dissolved in PBS to result in a final concentration of 25 uM. Immediately before the measurement, CoQ1-coumarin (Cayman Chemicals, 29554) in dissolved in PBS was added to the well and reduction of CoQ1-coumarin was observed on a TECAN Spark Plate reader with monochromator excitation of 405/20 nm and monochromator emission readout at 475/10 nm at a gain of 95. Data was plotted and analyzed with Graphpad Prism (10.2.2).

#### Artificial lipid droplet (aLD) synthesis

Artificial lipid droplets (aLD) were synthesized as previously published.^70^ Briefly, 10 umol of TG 18:1/18:1/18:1 (Nu-Chek Prep, T-235), 0.95 umol egg-PC (Avanti Research, 840051) and 18:1 DGS-NTA(Ni) (Avanti, 790404P) were each dissolved in 100 ul chloroform/methanol (2:1, v,v) and solutions were mixed. This resulted in a ratio of TG : egg-PC : 18:1 DGS-NTA(Ni) = 10 : 0.95 : 0.05. Lipid mixture was dried *in vacuo* in HPLC glass vials (Eppendorf, vaccuum concentrator 5301).

Dried lipids were resuspended in 1.25 ml ethanol by vigorous vertexing for 60s. To form artificial lipid droplets, 1.25 ml of lipids dissolved in ethanol were added dropwise to a stirring solution of 3.75 ml PBS containing 0.2% tween-20 (Sigma Aldrich, P7949). This solution was stirred for 4 hours at room temperature in an open 10 ml glass beaker until all ethanol had evaporated resulting in a solution containing 3.7 mM lipid.

#### Analysis of supramolecular structure of aLDs and liposomes via Nile Red fluorescence

aLDs or liposomes were diluted to 100 µM in 100 µl PBS and 2.5 µl of 250 µM solution of Nile Red (Sigma Aldrich, 19123) in ethanol were added and incubated for 15 min at room temperature. A TECAN Spark plate reader was used to read out Nile Red fluorescence by using monochromator excitation 530/10 nm with a gain of 100.

#### Quantification of aLD size and polydispersity by dynamic light scattering

Dynamic light scattering analysis was performed on a Brookhaven 90Plus Nanoparticle Analyzer. PBS was filtered three times through a 0.2 µm syringe filter (Pall Corporation, 4602) to remove particulate matter. A 3.5 mM stock solution of aLDs were diluted in filtered PBS 1:10,000 and analyzed.

#### Recruitment analysis of recombinant FSP1 to aLD

To probe successful recruitment of FSP1 to aLD, 2.5 mM aLD were incubated with 250 µM FSP1 (aLD : FSP1 = 1000 : 1) in a total volume of 250 µl for 1 hour at room temperature. aLD/FSP1 preparations were centrifuged at 17,000xg for 5 min at room temperature to float aLDs. The lower 175 µl were removed (unbound fraction) and the floating fraction was mixed with 175 µl PBS. aLD/FSP1 preparations were centrifuged again at 17,000xg for 5 min at room temperature to float aLDs. The lower 175 µl were collected (wash fraction) and 225 µl PBS were added to the remaining liquid (aLD bound fraction). 50 µl of each sample were mixed with SDS Page Laemmli buffer (Bio-Rad, 1610747) and separated via gel electrophoresis (Bio-Rad, 4561094). Bands were visualized via silver staining (Thermo Fisher, 24600) and band densities were quantified via densitometry.

#### Liposome synthesis

Liposomes were generated by mixing 16.2 µmol Egg-PC (Avanti Research, 840051) dissolved in chloroform/methanol (2:1, v/v) with 0.81 µmol 18:1 DGS-NTA(Ni) (Avanti, 790404P) dissolved in chloroform/methanol (2:1, v/v) in a HPLC glass vial followed by drying *in vacuo* (Eppendorf, vaccuum concentrator 5301). To form liposomes 324 µl PBS were added to dried lipids and vortexed for 60 seconds at room temperature yielding a final concentration of 50 mM lipid. Unilamellar liposomes were generated by extrusion using a mini-extruder (Avanti, 610000) by 19x passes through a 0.1 µm membrane (Cytiva, 230300).

#### Recruitment analysis of recombinant FSP1 to liposomes

To probe successful recruitment of FSP1 to liposomes, 10 mM liposomes were incubated with 1 µM FSP1 (liposomes : FSP1 = 1000 : 1) in a total volume of 100 µl for 45 minutes at room temperature. In ultracentrifugation tubes (Beckman Coulter, 3437755) 30 µl of liposome/FSP1 solution were mixed with 30 µl of 60% sucrose in PBS yielding a final concentration of 30% sucrose. This was overlayed with 100 µl 25% sucrose in PBS and subsequently with 20 µl PBS. Tubes were centrifuged at 100,000xg for 1h at 4C with a TLA.100 rotor on a Beckmann Coulter Optima TL Ultracentrifuge. Using gel loading tips, 155 µl of the bottom fraction were collected and discarded. The remaining floating fraction was mixed and collected. 16 µl of the floating fraction were mixed with SDS Page Laemmli buffer (Bio-Rad, 1610747) following vendor’s recommendation and separated via gel electrophoresis (Bio-Rad, 4561094). Bands were visualized via silver staining (Thermo Fisher, 24600).

#### Fluorescence enabled inhibited autooxidation (FENIX) assay in aLD and liposomes

BODIPY-C11 (Thermo Fisher Scientific, D3861) stock solutions were prepared in ethanol and stored at -80C until use. aLD or liposomes preparations were diluted to 2.5 mM and BODIPY-C11 was added to a final concentration of 2.5 µM (aLD : BODIPY-C11 = 1000 : 1). 40 µl of aLD/BODIPY-C11 or liposome/BODIPY-C11 solutions was added to a well in a 96 well plate followed by 1 µl of a solution of Coenzyme Q10 (Sigma Aldrich, C9538) or other FSP1 substrate dissolved in ethanol. To this solution was added 10 µl of a FSP1 solution,10 µl of a NADH solution and 40 µl PBS to achieve the desired concentration in a final reaction volume of 100 µl. This results in a final concentration of aLD/liposome (1mM) : BODIPY-C11 (1 µM) : CoQ10 : NADH. The oxidation initiator DTUN (Cayman Chemical, 32742) was dissolved in ethanol to yield a stock solution of 13.5 mM and 1.5 µl of DTUN stock were added to 100 µl reaction volume in well.

A TECAN Spark plate reader was preheated to 37C and plates were shaken in the plate reader for 60s using the “orbital” shaking mode with an amplitude of 5. Reduced BODIPY-C11 was detected by fluorescence through monochromator excitation at 488/20 nm and monochromator emission was detected at 590/10 nm at a gain of 120. Oxidized BODIPY-C11 was detected by fluorescence through excitation at 488/5 nm using a monochromator and emission was detected at 520/10 nm at a gain of 120.

#### DepMap analysis

Gene essentiality correlation with lipid abundance across cell lines was performed on the depmap portal (https://depmap.org/portal) as published before.^71^ A “custom analysis” was set up and “Pearson correlation” of the “gene = AIFM2 (FSP1, FLJ14497, AMID, PRG3)” from the “CRISPR (DepMap Public 24Q2+Score, Chronos) as “data slice” with the dataset “Metabolomics” of “all cell lines” was calculated. The generated data was downloaded as .csv file and plotted in GraphPad Prism 10.2.2 (GraphPad Software).

## Materials availability

All unique/stable reagents generated in this study are available from the lead contact with a completed Materials Transfer Agreement.

## Data availability

Raw data will be reposited on MASSive in the GNPS ecosystem for download.

## EXTENDED DATA FIGURE LEGENDS

**Extended Data Figure S1.**
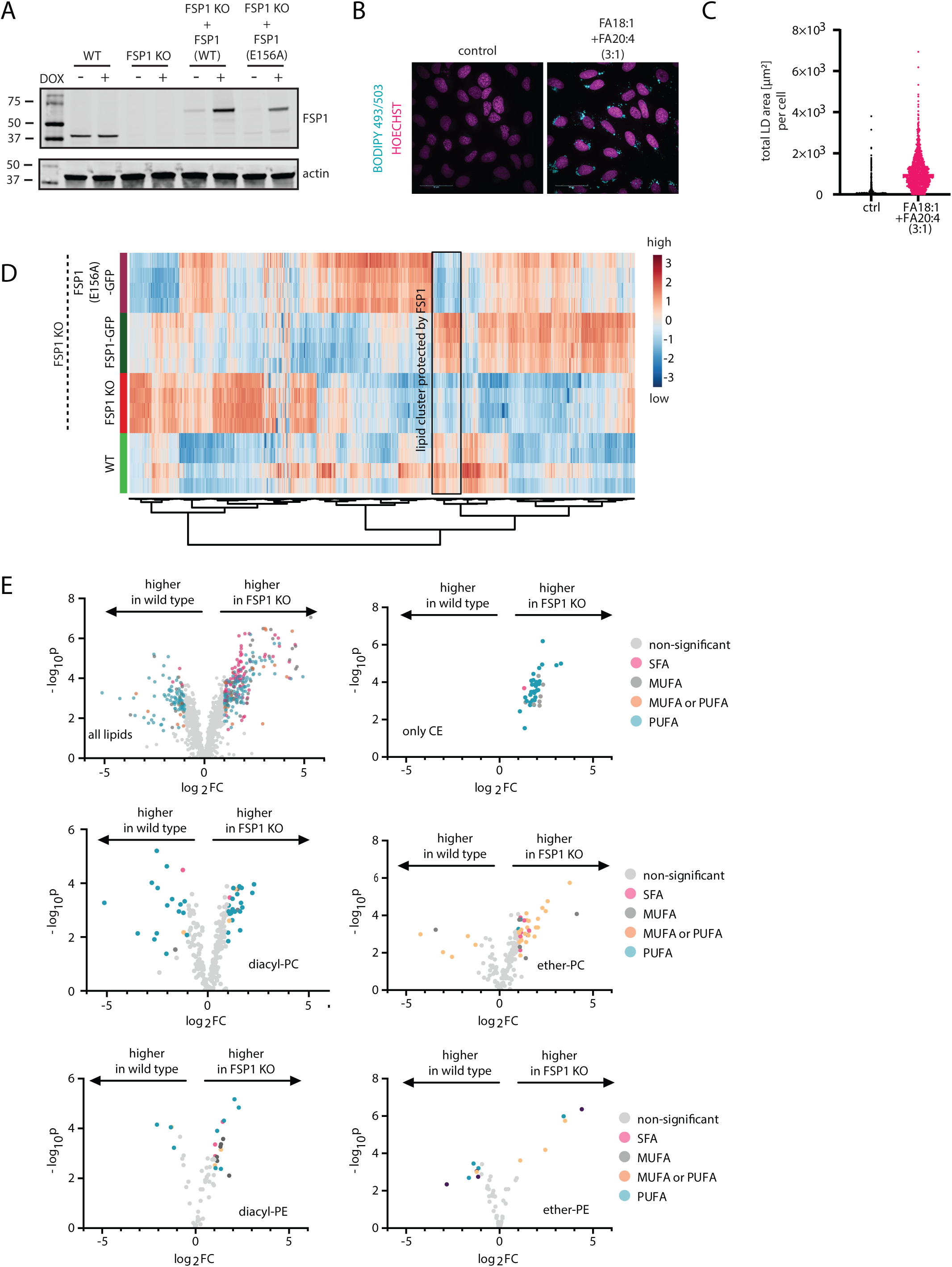
Loss of FSP1 result in lipid remodeling and a reduction in PUFA-containing glycerolipids. **A:** Western blot to assess FSP1 expression levels in used cell lines. DOX = doxycycline). **B:** Volcano plots of untargeted lipidomics data comparing all lipids, only cholesteryl esters, only diacyl-phosphatidylcholine, only ether-phosphatidylcholine (O and P linked), diacyl-phosphtadiylethanolamines, or ether-phosphatdiylethanolamines (O and P linked) of wild type vs FSP1 KO cells. Triacylglycerol unsaturation is colored according to legend. SFA = saturated fatty acid containing (total double bonds = 0), MUFA = monounsaturated fatty acid containing (total double bonds = 1), PUFA = polyunsaturated fatty acid containing (total double bonds ≥ 4), MUFA or PUFA (total double bonds = 2 or 3). (data was autoscaled and log10 transformed, thresholds for significant regulation are fold change > 2, p < 0.05 (FDR corrected)). **C:** Lipid droplet imaging of wild type cells treated with oleate (FA 18:1) and arachidonate (FA 20:4). Lipid droplets were stained with BODIPY 493/503 and nuclei with Hoechst. **D:** Lipid droplets were imaged and lipid droplet area for each cell was quantified using Harmony data analysis software (Revvity). Each dot represents the lipid droplet area in an individual cell. **E:** Heatmap displaying all lipids quantified in each cell line. Hierarchical clustering was performed using Metaboanalyst 5.0 and a cluster of lipids specifically protected by FSP1 is indicated by the black box. (data was autoscaled and log10 transformed)

**Extended Data Figure S2.**
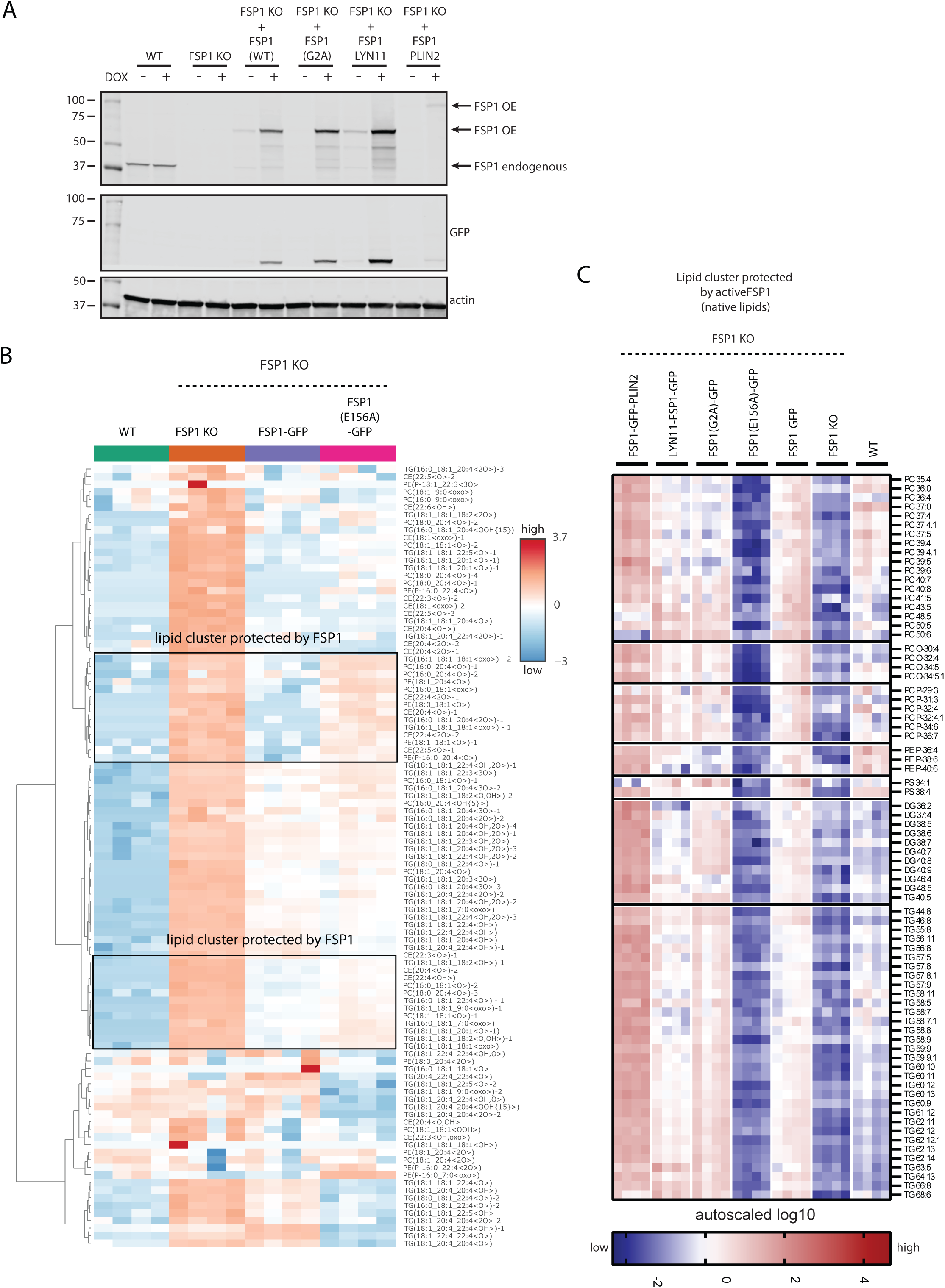
Lipid droplet-localized FSP1 prevents neutral lipid peroxidation. **A:** Western blot to assess FSP1 expression levels in used cell lines. DOX = doxycycline. **B:** Abundances of all oxidized lipids identified and quantified within different FSP1 expression constructs. Hierarchical clustering was performed using Metaboanalyst 5.0 and a cluster of lipids specifically protected by FSP1 is indicated by the black box (data was autoscaled and log10 transformed). **C:** Heatmap displaying lipid levels within different localization mutants from a cluster of lipids that is protected by catalytically active FSP1 as identified by hierarchical clustering in Figure S1E (data was autoscaled and log10 transformed).

**Extended Data Figure S3.**
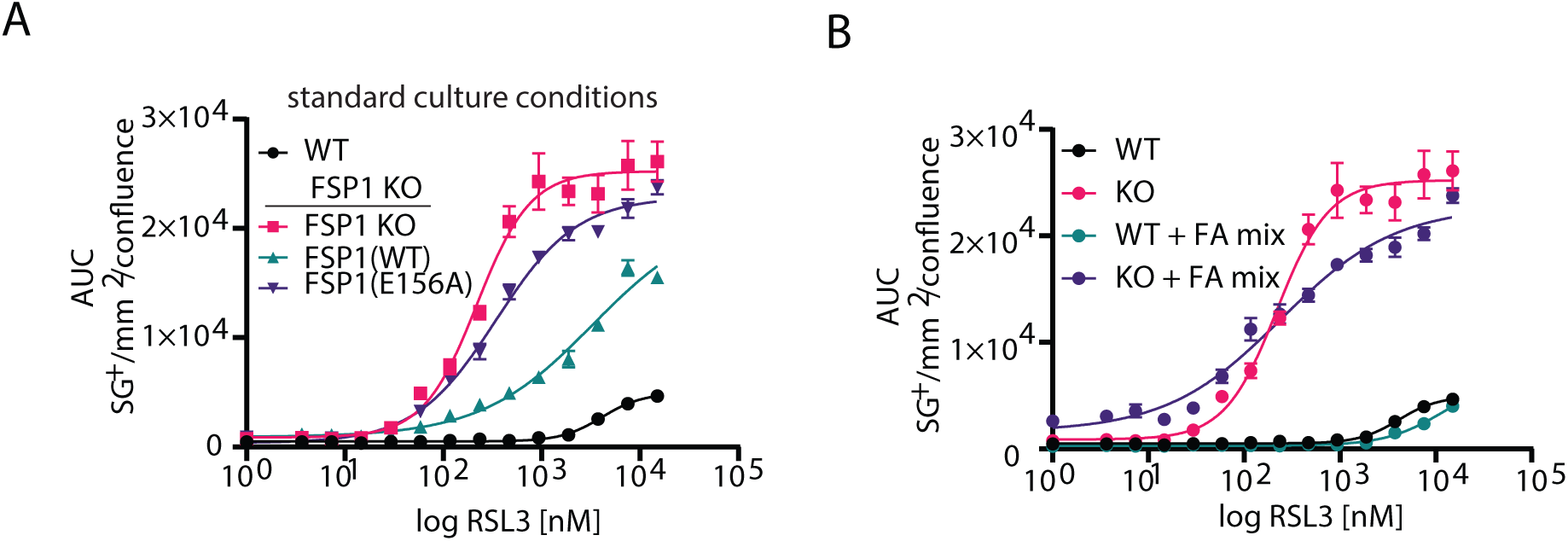
Lipid droplet-localized FSP1 prevents ferroptosis. **A,B:** Cell death sensitivity to RSL3 assessed by time resolved fluorescence microscopy (incucyte) with cell death indicator sytox green over the course of 24 hours. Number of sytox green positive objects per image equals the number of dead cells and is normalized to cell area per image expressed as confluency. sg+ = sytox green positive objects.

**Extended Data Figure S4.**
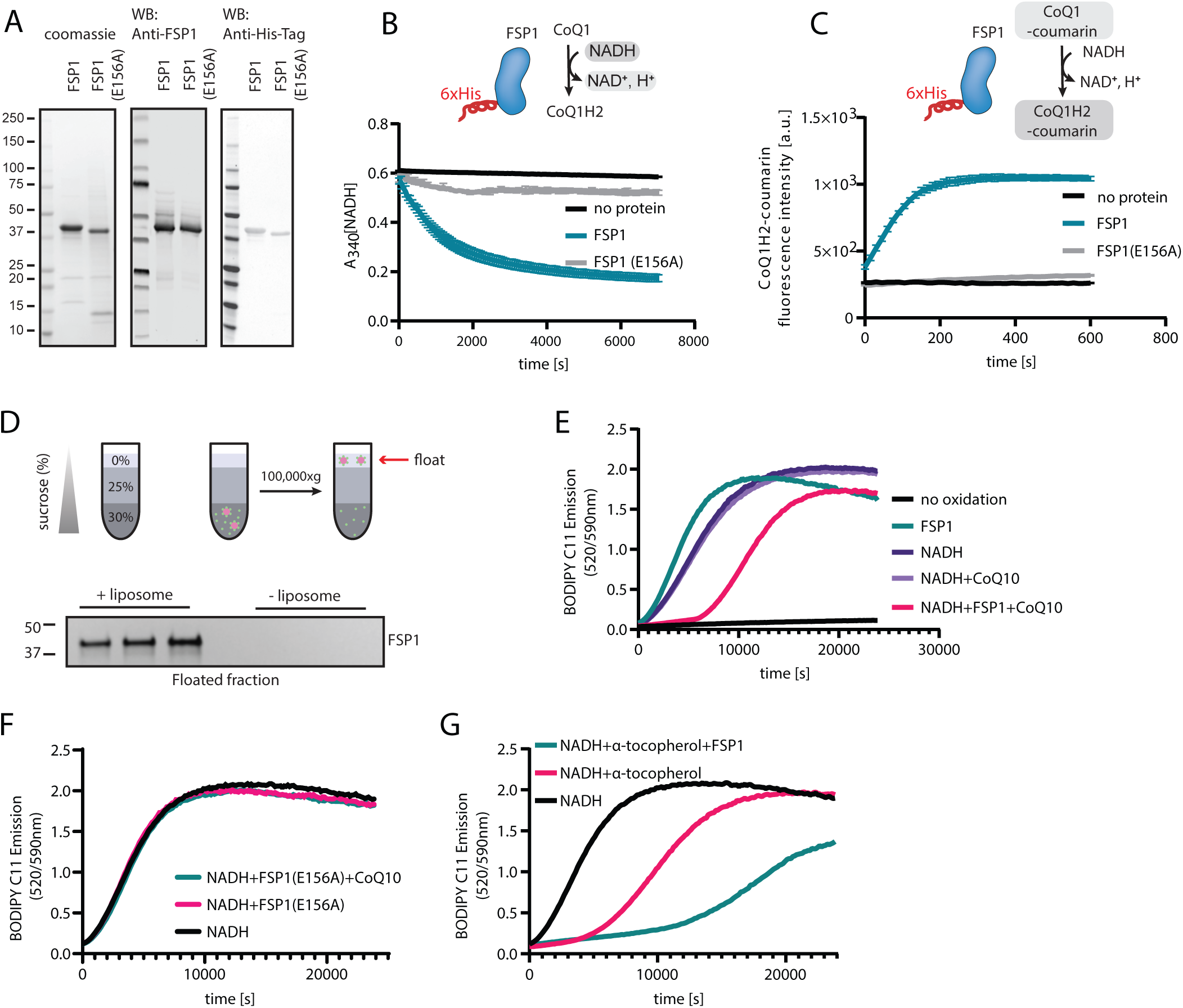
Recombinant FSP1 reduces CoQ10 by consuming NADH to suppress phospholipid peroxidation in liposomes in vitro. **A:** Characterization of recombinantly expressed FSP1. Coomassie stained gel indicates high FSP1/FSP1(E156A) purity and western blot analysis proves the identity of FSP1 and the presence of a His-Tag. **B:** Recombinant FSP1 but not FSP1(E156A) is capable of consuming NADH in the presence of the soluble CoQ10 derivative CoQ1. NADH consumption is assessed by measuring endogenous absorbance of NADH at 340 nm. **C:** Recombinant FSP1 but not FSP1(E156A) can reduce CoQ to CoQH2 in the presence of NADH. CoQH2 formation is assessed using the fluorescent CoQ-reduction reporter CoQ1-coumarin. **D:** Recombinant FSP1 can be recruited to nickel-phospholipid doped liposomes. Recruitment was assessed using a liposome flotation assay and gel electrophoresis with silver stain detection. **E:** FENIX assay is Egg-phosphatidylcholine liposomes in the presence of various substrate and enzyme combinations. **F:** FENIX assay in Egg-phosphatidylcholine liposomes in the presence of catalytically inactive FSP1(E156A). **G:** FENIX assay in Egg-phosphatidylcholine liposomes in the presence of α-tocopherol and/or FSP1.

**Extended Data Figure S5.**
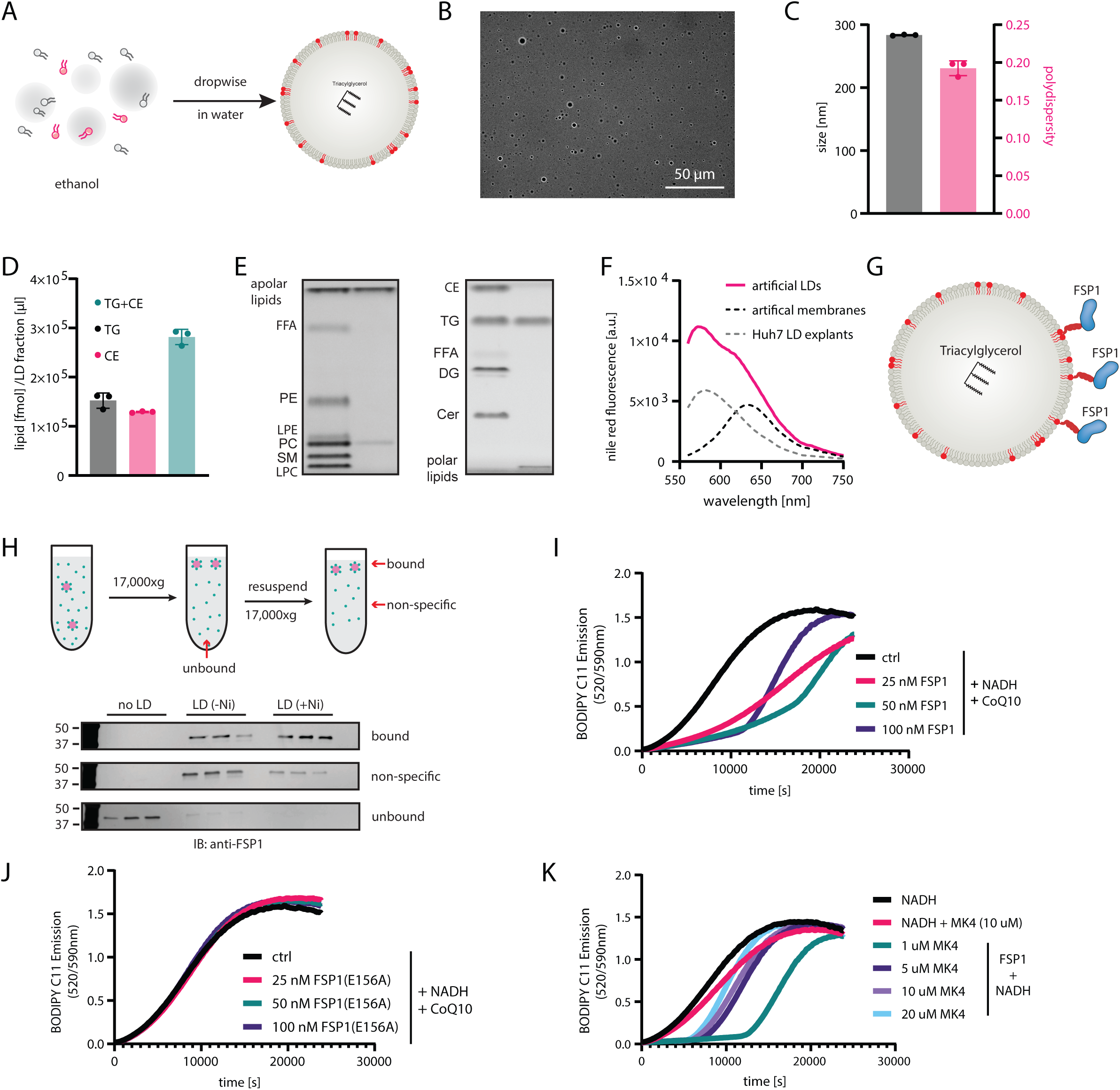
Artificial lipid droplets recapitulate cellular lipid droplets and recruitment of recombinant FSP1 prevents lipid peroxidation in vitro. **A:** Schematic of ethanolic infusion of lipids to synthesize artificial lipid droplets. **B:** Phase contrast microscopy of artificial lipid droplet preparations. **C:** Size and polydispersity of artificial lipid droplet preparations assessed by dynamic light scattering. **D:** Lipid class distribution in lipid droplet fractions extracted from U-2 OS cells treated with 200 uM oleate and 0.1% BSA for 24 hours. Lipids were quantified by thin layer chromatography. **E:** Thin layer chromatograms of lipids artificial lipid droplets. **F:** Nile red fluorescence in liposomes, artificial lipid droplets and lipid droplets extracted from Huh7 cells. Nile red was excited at 530/10 nm and shows red-shifted fluorescence in comparison to liposomes but similar fluorescence as compared to lipid droplet fractions from cells indicating a lipid droplet like supramolecular structure. **G:** Schematic describing the strategy to recruit His-tagged FSP1 to nickel-phospholipid spiked artificial lipid droplets. Red phospholipids = DGS-NTA(Ni), red tail on FSP1 = His-Tag. **H:** Schematic describing artificial lipid droplet flotation to probe successful recruitment of His-tagged FSP1. Lower panel shows silver stained gel detecting FSP1 protein in each collected fraction indicating strong binding. **I:** FENIX assay in artificial lipid droplets in the presence of 100 nM FSP1, 25 µM NADH and varying concentrations of the alternative FSP1 substrate menaquinone 4 (MK4). **J:** FENIX assay in artificial lipid droplets in the presence of 10 µM CoQ10, 25 µM NADH and varying concentrations of the catalytically dead FSP1 mutant FSP1(E156A). **K:** FENIX assay in artificial lipid droplets in the presence of 10 µM CoQ10, 25 µM NADH and varying concentrations of active FSP1 mutant FSP1.

**Extended Data Figure S6.**
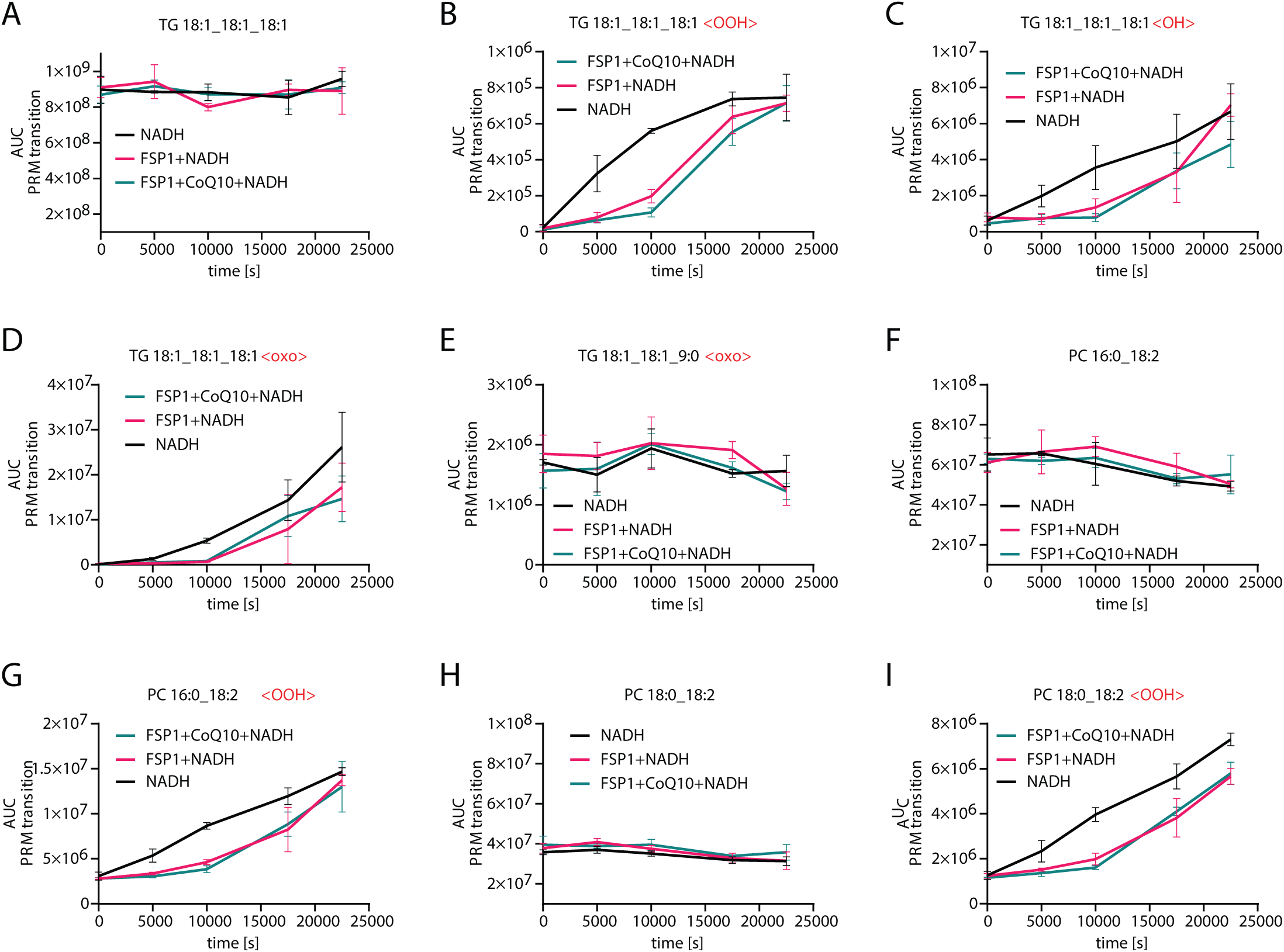
In vitro peroxidation of artificial lipid droplets is characterized by phospholipid and triacylglycerol peroxidation and FSP1 prevents lipid peroxidation in a CoQ and NADH dependent manner. **A-I:** Quantification of lipid peroxidation products detected by parallel reaction monitoring during peroxidation of artificial lipid droplets in the presence of 25 uM NADH, 10 uM CoQ10 and 100 nM FSP1.

**Extended Data Figure S7.**
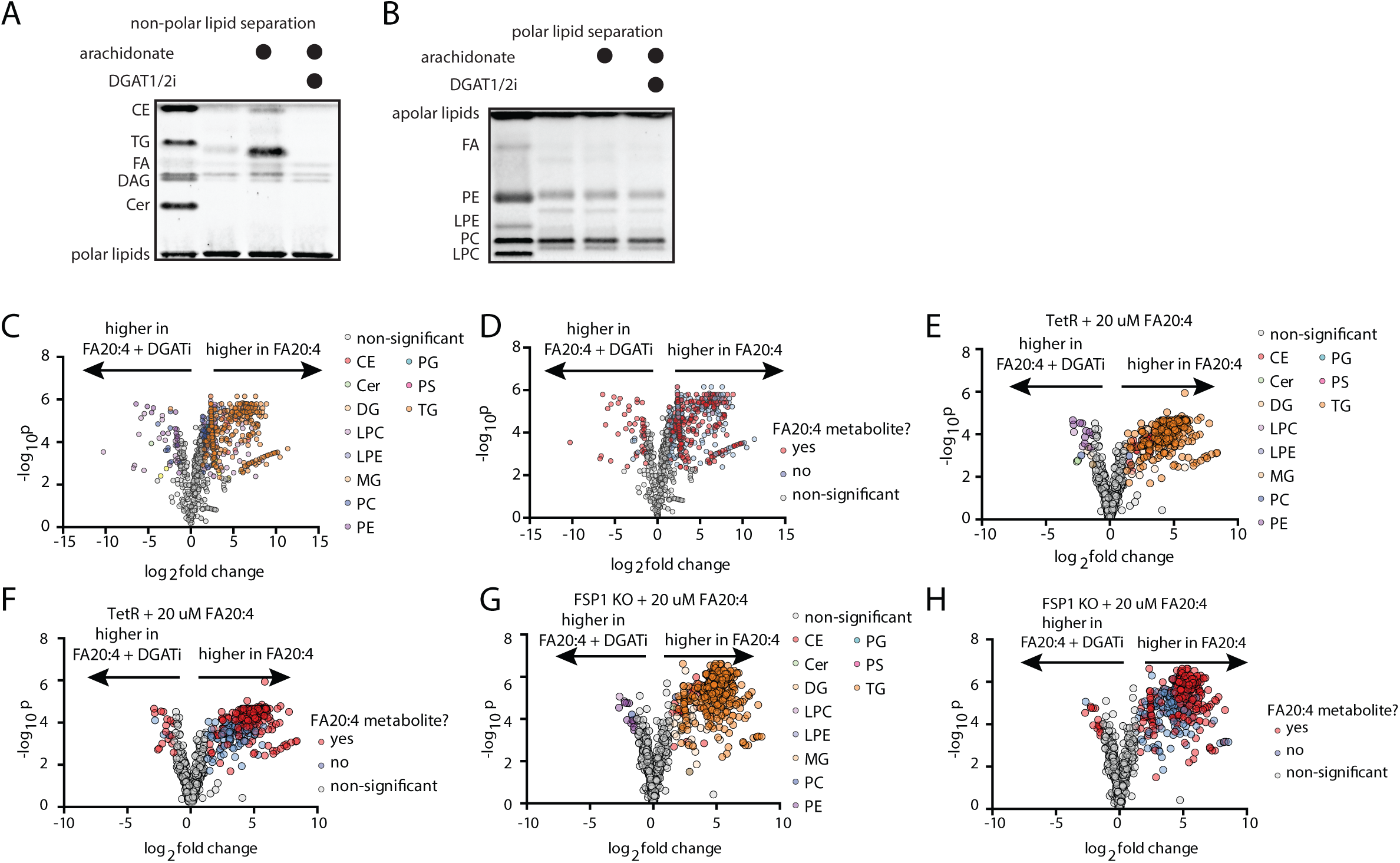
Arachidonic acid treatment induces specific accumulation of polyunsaturated non-polar lipids. **A,B:** Lipid extracts from cells treated with 200 uM FA20:4 for 24 hours in the presence and absence of DGAT1/2 inhibitors (15 μM A-922500 and 10 μM PF-06424439). Non-polar and polar lipids are separated and quantified by thin layer chromatography. **C,E,G:** LC-MS/MS based untargeted lipidomics of conditions described in the graph titles. Lipid classes are colored according to legend. Cer = ceramide, DG = diacylglcyerol, TG = triacylglycerol, CE = cholesteryl ester, PC = phosphatidylcholine, LPC = lysophosphatidylcholine, PE = phosphatidylethanolamine, LPE = lysophosphatidylethanolamine, SM = sphingomyelin, PS = phosphatidylserine, PI = phosphatidylinositol, PG = phosphatidylglycerol. (data was autoscaled and log10 transformed, thresholds for significant regulation are fold change > 2, p < 0.05 (FDR corrected)). **D,F,H:** Same data as displayed in Fig S6A/C/E. Significantly regulated lipids are colored based on the presence or absence of FA20:4 metabolites. A lipid was deemed a putative FA20:4 metabolite when either of its fatty acids had more than 20 carbons with more than 4 double bonds.

**Extended Data Figure S8.**
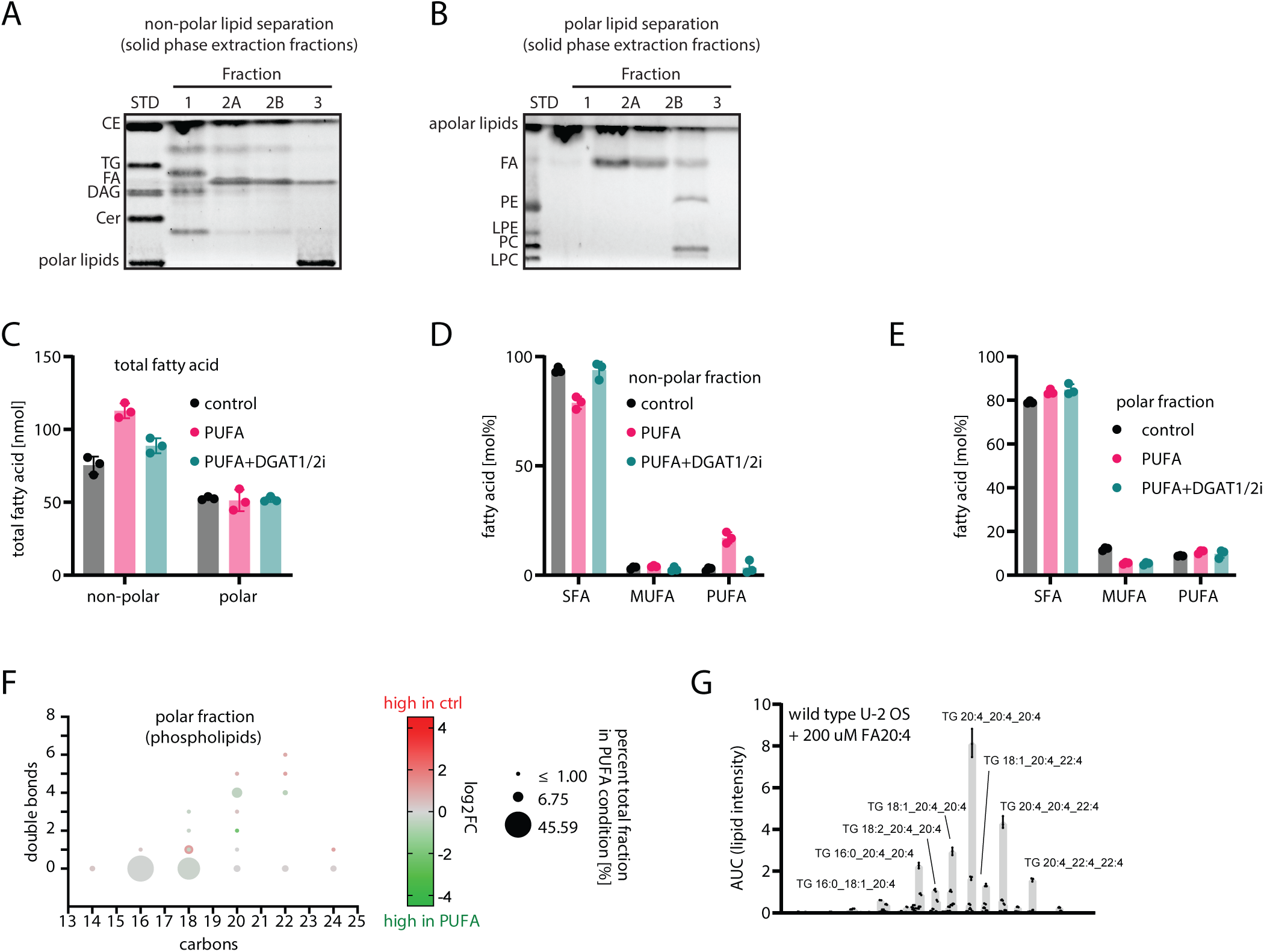
Quantitative analysis of fatty acid composition following arachidonic acid treatment. **I/J:** Lipid extracts from cells were separated into fractions and non-polar and polar lipids were separated by thin layer chromatography. **K:** Lipid extracts were separated, and esterified fatty acids were quantified. All fatty acid concentrations were summed and represent total fatty acid concentrations in each fraction. **L:** Concentrations of saturated fatty acids (SFA), monounsaturated fatty acids (MUFA) and polyunsaturated fatty acids (PUFA) in each fraction were summed up. **N: J:** Bubble plot showing absolute quantities and regulation of fatty acids in the polar lipid fraction extracted from U-2 OS cells treated with 200 µM FA20:4 for 24 hours. For quantification, lipid extracts were separated into non-polar lipid fraction using solid phase extraction and esterified fatty acids were transmethylated and quantified using gas chromatography-flame ionization detection. **O:** Triacylglycerol lipid intensity from untargeted lipidomics indicating the simplicity of the triacylglycerol species under the studied conditions.

**Extended Data Figure S9.**
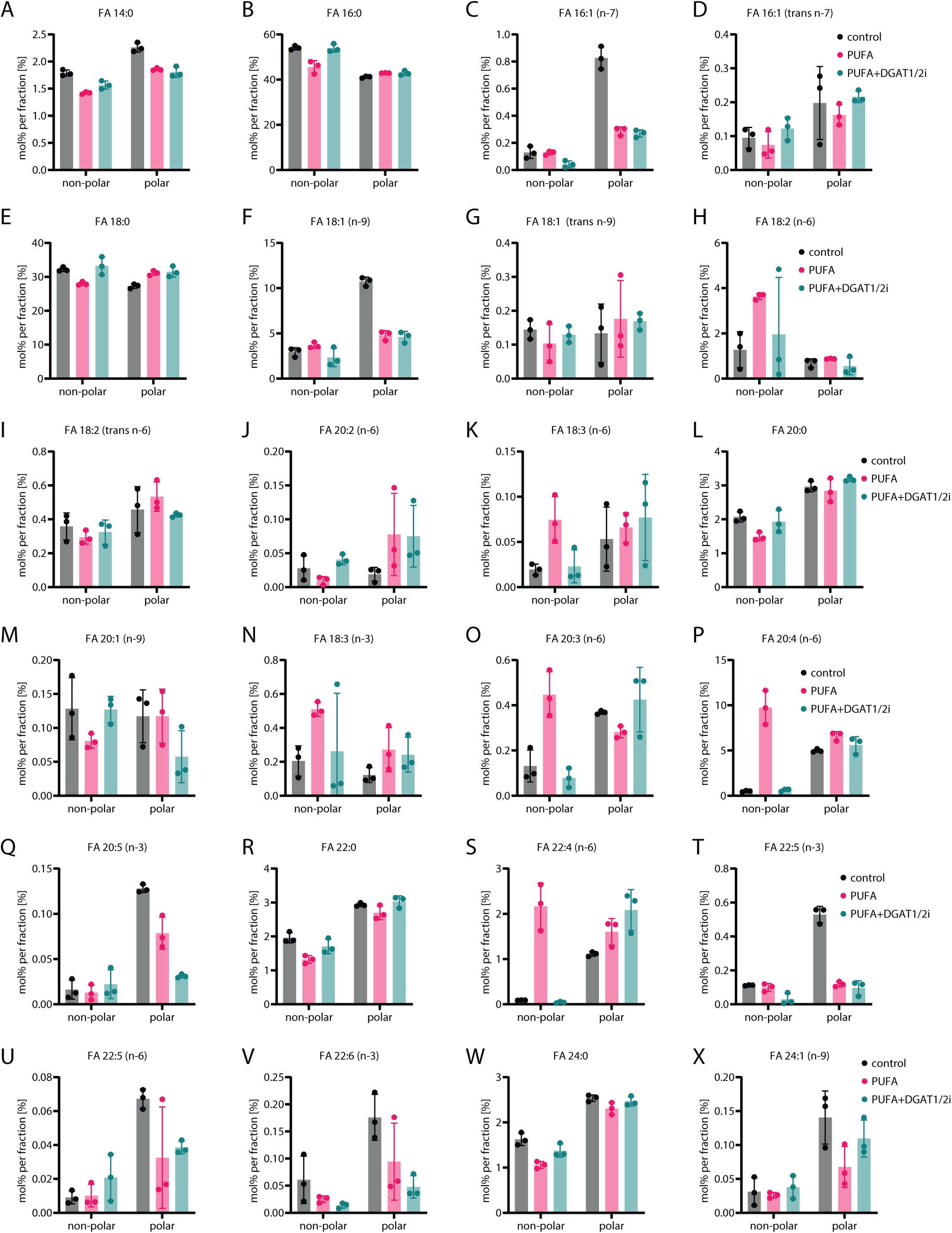
Arachidonic acid treatment induces specific accumulation of arachidonic acid containing non-polar lipids and is blocked by DGAT1/2 inhibition. **A-X**: Cells were treated with 200 µM FA20:4 for 24 hours in the presence and absence of DGAT1/2 inhibitors (15 μM A-922500 and 10 μM PF-06424439). Lipids were extracted from cell pellets. Lipid extracts were separated into polar and non-polar lipids using solid phase extraction. Esterified fatty acids were released and transmethylated. Fatty acid methyl esters were absolutely quantified using gas chromatography coupled to flame ionization detection.

**Extended Data Figure S10.**
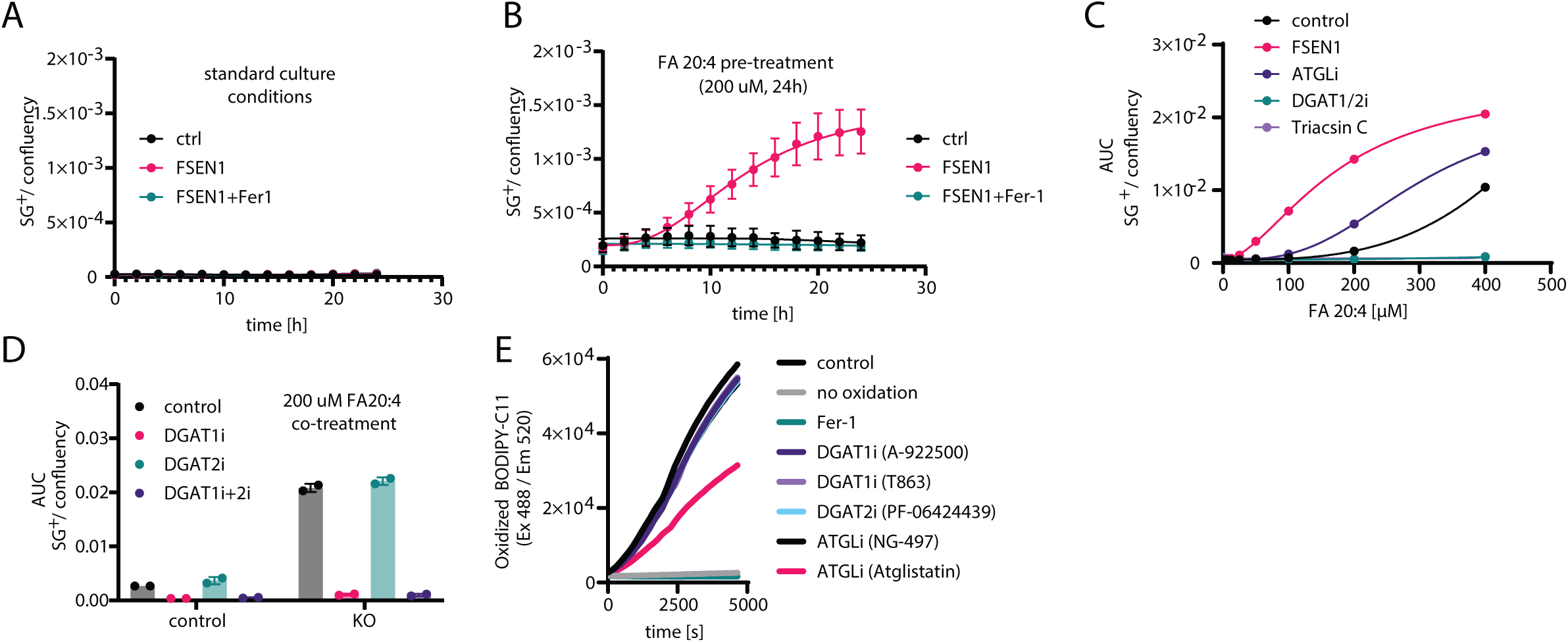
Non-polar lipid regulating compounds regulated lipid droplet initiated Ferroptosis and not by an off-target activity of lipid metabolism inhibitors. **A-D** : Cell death assess by time resolved fluorescence microscopy (incucyte) with cell death indicator sytox green. **E:** Fluorescence enabled inhibited autooxidation in Egg-PC liposomes detecting no radical trapping activity of utilized lipid metabolism inhibitors.

